# Determinants of Blood Group Antigen Expression and Prediction of Phenotypes by Machine Learning

**DOI:** 10.64898/2026.07.01.735824

**Authors:** Anne-Cathérine Kranz, Johannes Schneider, Christoph Gassner, Maike Bublitz

## Abstract

**Key Points:**

- Structural and evolutionary features distinguish null and antigenic blood group variants across multiple systems.
- Machine learning can predict variant effects, enabling improved interpretation of novel mutations in transfusion medicine.

Blood group antigens, defined by epitopes on the erythrocyte surface, are central to transfusion safety and maternal–fetal compatibility. While the genetic basis of many clinically relevant blood group antigens is well established, which structural and biophysical parameters determine whether a single-nucleotide variant gives rise to an antigenic phenotype remains unclear. Here, we integrate structural, biophysical, and evolutionary analyses to systematically evaluate features associated with single amino acid substitutions across 24 human protein-based blood group systems. We analyse 319 variants with curated phenotypic annotations alongside 481 control variants, identifying key determinants of null and antigenic phenotypes. Null variants are characterized by high evolutionary conservation, burial within the protein core, loss of hydrophobicity, increased polarity, and a propensity for arginine substitutions. Antigenic variants are also enriched in arginine; however, in contrast to null variants, they tend to occur at less conserved, more solvent-accessible, and structurally flexible sites. Supervised machine learning models trained on structural and biophysical descriptors were applied to distinguish (i) null and (ii) antigenic variants from controls, achieving balanced accuracies of 0.82 and 0.63, respectively. Feature importance analysis identified predicted pathogenicity, solvent accessibility, and evolutionary conservation as the most predictive determinants of null variants, whereas hydrophobicity, conservation, and flexibility dominated antigen prediction. This work establishes a framework linking molecular variation to blood group phenotypes and provides a foundation for predicting the impact of novel missense mutations in transfusion medicine and beyond.

## 1 Introduction

Red blood cell antigens are genetically encoded protein or carbohydrate markers that play a crucial role in transfusion safety and maternal-fetal health. One or multiple related antigens encoded by a single gene or closely linked genes constitute a blood group system^1^. As of October 2025, 48 human blood group systems are recognized by the International Society of Blood Transfusion (ISBT) Red Cell Immunogenetics and Blood Group Terminology Working Party^2,3^. Incompatible transfusions can provoke haemolytic transfusion reactions, and haemolytic disease of the foetus and newborn remains an important cause of stillbirth and neonatal death in cases of maternal-foetal incompatibility^4^.

Blood group antigens are determined by specific alleles, typically representing defined haplotypes across the respective gene. The reference allele defines the canonical antigen(s). Genetic variation ranges from single nucleotide variants (SNVs), which can cause amino acid substitutions, to recombination events (including gene conversions), inversions, insertions, and deletions^5^. Serological antibody identification remains indispensable in routine practice to prevent adverse transfusion outcomes. In parallel, molecular high-throughput approaches, ranging from SNV genotyping to sequencing, are increasingly applied to predict blood group phenotypes and refine donor-recipient matching. Beyond transfusion medicine such predictions are relevant to fields such as autoimmunity and cancer. The vast case numbers in blood donor genotyping and serological antigen detection makes blood group proteins a formidable model system for computer-aided prediction of clinical phenotypes. Furthermore, the pool of known non-rare polymorphisms in the European population represents a valuable control group, as clinical phenotypes can be ruled out with high confidence for these variants, given the high frequency of testing.

Machine learning has been successfully applied to blood group antigen determination from genomic data^6^, and efforts to predict antigenicity have expanded: Approaches include mathematical models for estimating immunogenicity and/or alloimunization risk from medical data such as antigen and antibody frequencies, transfusion counts, antigen density and recipients’ responsiveness^7,8^. Protein structural modelling combined with epitope prediction and/or molecular dynamics simulations have been used to describe antigens or variants in individual blood group systems^9–16^.

Recent studies suggest that structure-based and biophysical parameters such as solvent accessibility and flexibility, can correlate with blood group phenotypes^13,17^. These analyses focused on individual blood group systems or small antigen sets. Here, we expand this concept into a comprehensive structural characterization and prediction of effects of single-amino acid variants across multiple blood group systems. We asked whether measurable, significant features distinguish antigenic or null phenotypes — i.e., missense variants that abolish antigen expression without truncating the protein — across systems and whether such features can train machine learning (ML) algorithms to identify phenotypes. To address this, we compiled all known single–amino acid substitutions with ISBT curated phenotypic annotations across 24 blood group systems, calculating seven structural, biophysical and conservation-based descriptors for 800 missense variants. We then compared feature distributions and trained ML models to predict antigenic and null phenotypes. Feature importance analysis identified key molecular determinants underlying antigenicity and loss of expression.

## 2 Materials and Methods

### 2.1 Data sources

The 24 blood group systems included in this study and the number of single missense variants with known phenotypes (excluding frame shifts or truncations) are shown in **Table 1**.

**Table 1:** Number of blood group single amino acid variants used in this study. Variants are classified as null variants (NullV), defined by complete loss of expression of all antigens encoded by a specific blood group gene; control variants (CtrlV), representing amino acid substitutions that do not result in antigen expression; and antigenic variants, subdivided into wild-type antigens (AgWT) and non-wild-type antigenic forms (AgV), including antithetical variants (relative to AgWT) and de novo variants.

| ISBT-Number | Blood group<br>system | CtrlV | AgWT | AgV | NullV | Total Count |
| --- | --- | --- | --- | --- | --- | --- |
|  |  | No<br>phenotype | Antigenic<br>wild-type | Antigenic<br>variant | Null<br>phenotype |  |
| 002 | MNS (GYPA) | 10 | 8 | 17 | 0 | 35 |
| 002 | MNS (GYPB) | 10 | 2 | 12 | 0 | 24 |
| 004 | RhD | 46 | 37 | 0 | 3 | 86 |
| 005 | Lutheran | 60 | 24 | 4 | 0 | 88 |
| 006 | Kell | 38 | 23 | 14 | 18 | 93 |
| 008 | Duffy | 16 | 1 | 2 | 4 | 23 |
| 009 | Kidd | 10 | 1 | 1 | 21 | 33 |
| 010 | Diego | 48 | 3 | 21 | 1 | 73 |
| 011 | YT | 22 | 4 | 1 | 0 | 27 |
| 013 | Scianna | 22 | 6 | 2 | 0 | 30 |
| 014 | Dombrock | 12 | 8 | 1 | 1 | 22 |
| 015 | Colton | 11 | 1 | 3 | 1 | 16 |
| 016 | Landsteiner<br>Wiener | 11 | 1 | 1 | 0 | 13 |
| 020 | Gerbich (GYPC) | 13 | 5 | 4 | 0 | 22 |
| 021 | Cromer | 7 | 14 | 3 | 0 | 24 |
| 023 | Indian | 11 | 5 | 1 | 0 | 17 |
| 024 | Ok | 15 | 3 | 0 | 0 | 18 |
| 025 | Raph | 8 | 3 | 0 | 0 | 11 |
| 026 | John Milton<br>Hagen | 34 | 9 | 0 | 0 | 43 |
| 030 | RhAG | 13 | 2 | 3 | 10 | 28 |
| 035 | CD59 | 3 | 1 | 1 | 0 | 5 |
| 036 | Augustine | 22 | 2 | 1 | 0 | 25 |
| 037 | Kanno | 7 | 1 | 0 | 0 | 8 |
| 039 | CTL2 | 32 | 3 | 1 | 0 | 36 |
|  | **Total** | **481** | **167** | **93** | **59** | **800** |

Variant data were obtained from the ISBT Blood Group Allele Tables^18^ and the GnomAD database^19^. The dataset included antigens present in reference alleles denoted as antigenic wild-type residues (AgWT, n=167), antigenic variant residues (AgV, n=93) encoded by variant alleles, null variants (NullV, n=59) which are substitutions that lead to loss of detectable antigen expression, and control variants (CtrlV, n=481), which are non-rare variants with no known phenotype that were used as controls (defined as GnomAD search hits within the European non-Finnish population with a minor allele frequency (MAF) of >3.0 × 10^-5^).

Protein structures were retrieved from the Protein Data Bank (www.rcsb.org)^20^ or predicted using Alphafold2^21^ (**Supplemental Table 1**). Structures were manually curated to ensure physiological oligomers/context within protein complexes where possible. Amino acid variants were mapped onto 3D structures using residue numbering from reference sequences^22^.

For each variant position, seven structural, biophysical and conservation-based descriptors were calculated (**Table 2**). For null variant prediction, mutation pathogenicity values from AlphaMissense^23^ were included. Hydrophobicity and sidechain volume differences between the wild-type and variant residues were calculated to quantify physicochemical change. For a complete list of variants and all descriptors see **Supplemental Table 2**.

**Table 2:**
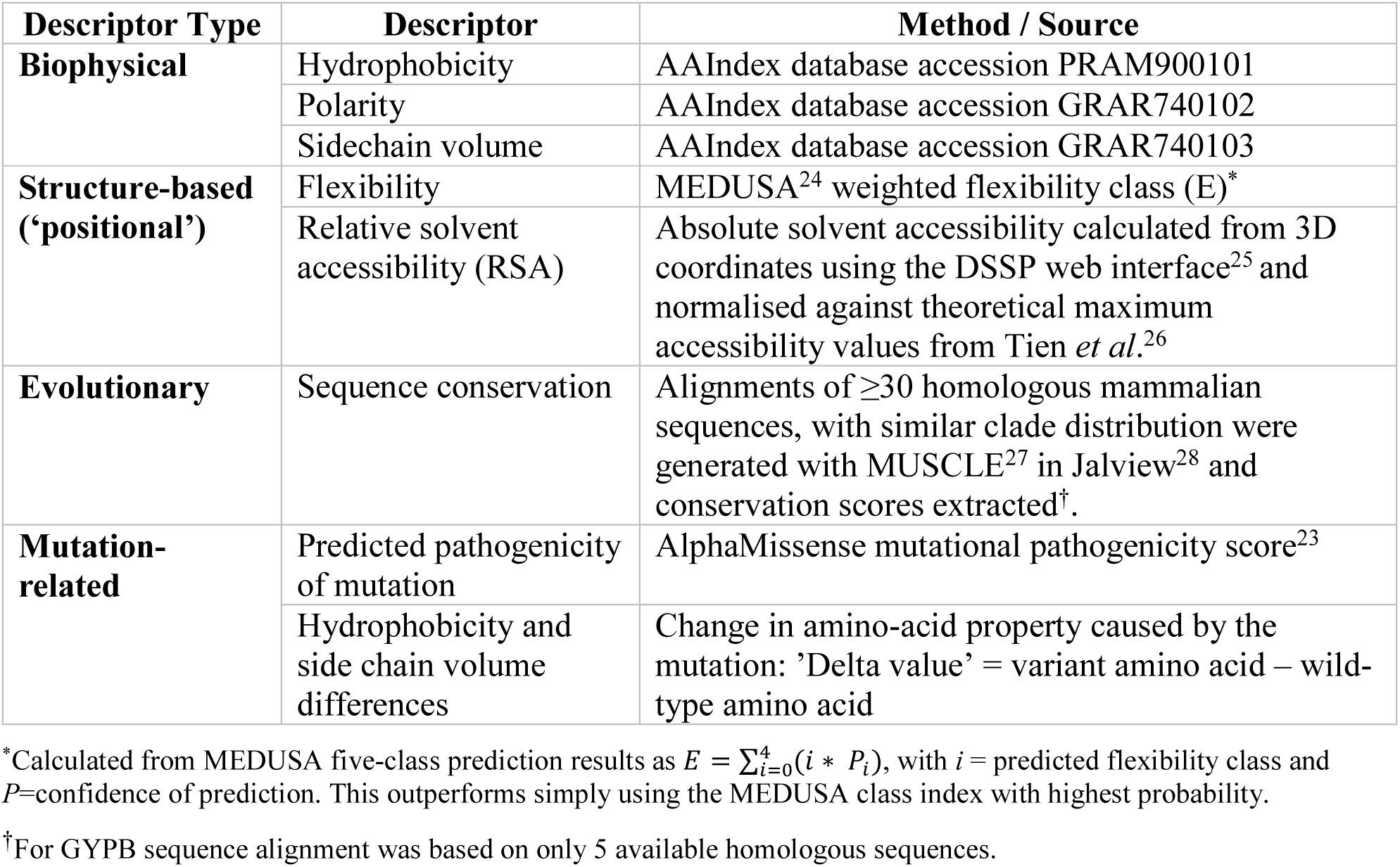
Set of descriptors calculated for each amino acid variant position.

### 2.2 Statistical Analysis

To compare the collected descriptor values across variants, we performed statistical analyses and visualized the distributions with violin plots. Summary statistics were overlaid on each group/half as median (short horizontal line or dot) and interquartile range (IQR) (short vertical line). Data points were displayed as x-jittered point overlays sorted by value (split violins only) to show sample density. For clarity, hydrophobicity values were sign-flipped in split-violin figures where indicated so that larger values consistently represent increased hydrophobicity.

### 2.3 Amino Acid Frequency Analysis

The frequencies of each amino acid within the groups NullV, AgWT+AgV and CtrlV were calculated by dividing the number of occurrences by the total number of variants in the group. These values were then normalised to the baseline frequencies of each amino acid within the full protein sequences of the 24 analysed blood group proteins, which was set to 1. This allowed a comparison of each amino acid’s occurrence in each group relative to its expected occurrence within the full set of proteins that were analysed. When comparing changes induced by mutations, amino acid residues before mutation were denoted ‘NullV-WT’ and ‘CtrlV-WT’, and after mutation ‘NullV-V’, and ‘CtrlV-V’.

### 2.4 Predicting null and antigenic variants by machine learning (ML)

Supervised ML models were employed to predict the impact of missense mutations on the antigenicity and expression of blood group protein variants. Two different classification tasks were performed: (i) Antigenic vs. Control and (ii) Null vs. Control. The models used were: Random Forest, Gradient Boosting, Support Vector Machine (SVM), and Logistic Regression. These classifiers were selected because they are well-established and well-performing models for small-to-medium tabular datasets. In addition, they represent linear probabilistic (Logistic Regression), linear maximum-margin (SVM), and tree-based (Random Forest, Gradient Boosting) modelling approaches, which reduces the reliance on a single set of assumptions and enables assessment whether identified signals are robust across model classes. Importantly, they support interpretability through transparent model structure or post-hoc explanations, which is critical in our biomedical prediction setting^29^.

To ensure a robust estimation of model performance, model training and evaluation were performed using group-aware, stratified nested cross-validation. The outer cross-validation loop consisted of 3 folds, while hyperparameter tuning was conducted within an inner 2-fold cross-validation loop to preserve both class distribution and group structure. Full search grids are provided as code in **Supplemental data**. Grouping was done at the blood group system level, to avoid information that makes the blood group system statistically dependent, (protein structure information and evolutionary background within a blood group) to leak from the training set to the test set. This allows generalization to unseen blood group systems and provides an unbiased estimate of model performance. In each outer fold, ∼67% of variants were used for training and ∼33% for testing. The reported performance values and feature-importance results correspond to a fixed outer cross-validation split configuration generated with random seed 42. Class imbalance was addressed by applying random undersampling of the majority class within the training folds only. Where supported, additional balanced weighting was used. Data pre-processing steps were: handling missing values by mean imputation, feature standardization with z-score scaling and encoding target class labels into numeric categories. Variants with mismatched wild-type residues between AlphaMissense annotations and the curated dataset (due to differences in sequences used by ISBT and AlphaMissense) were set to missing (NaN) prior to modeling, and subsequently handled by imputation within the cross-validation pipeline. To assess the performance, the following six metrics are reported: accuracy, weighted precision, weighted recall, weighted F1, Macro-F1 and balanced accuracy (BalAcc). BalAcc accounts for class imbalance by averaging sensitivity and specificity, while Macro-F1 reflects the harmonic mean of precision and recall computed independently for each class.

Feature importance was extracted separately within each outer cross-validation fold from the fitted classifiers. For Random Forest and Gradient Boosting, importance was defined using the models’ native impurity-based feature importance scores. For Logistic Regression and the linear SVM, importance was defined as the normalized absolute value of the fitted model coefficients. Importance values were normalized within each fold to sum to one, and are reported as mean ± standard deviation across outer folds.

Pairwise model comparisons were performed using generalized repeated grouped 5×3 paired tests on outer-fold performance differences, specifically Dietterich-like t-tests and Alpaydin-like F-tests, with Holm correction for multiple comparisons. For full details and code including multicollinearity assessment, variance inflation factor analyses, and an AlphaMissense ablation analysis see **Supplemental data.**

## 3 Results

We analysed single-amino-acid variants across multiple blood group systems, grouping them according to their antigenic effects: Null variants (NullV) were defined as substitutions leading to complete loss of expression of all antigens encoded by the respective blood group gene. Control variants (CtrlV) comprised amino acid substitutions without known phenotypic effects that have a minor allele frequency (MAF) of at least 3 × 10⁻⁵ in the European non-Finnish population as listed in gnomAD. Antigenic variants were comprised of wild-type antigens (AgWT) and non-wild-type antigenic forms (AgV), the latter including antithetical variants (relative to AgWT) as well as de novo variants.

### 3.1 Comparison of null variants with control variants

#### 3.1.1 Qualitative Analysis

We first compared the distributions of hydrophobicity, side chain volume and polarity between null variants (NullV), control variants (CtrlV), and their respective wild-type residues (**Error! Reference source not found.**).

NullV are associated with decreased hydrophobicity whereas CtrlV show distributions similar to wild-type residues (**Figure 1A**). Side chain volume increases more in NullV than in CtrlV (**Figure 1B**). Polarity increases in NullV and decreases in CtrlV (**Figure 1C**). Together, these shifts – reduced hydrophobicity, increased polarity and larger side chains – are consistent with destabilizing substitutions that disrupt the tightly packed hydrophobic protein core in null variants, impairing protein expression. In contrast, CtrlV variants resemble wild-type residues, indicating tolerated substitution without phenotypic effect. These findings support the hypothesis that disruptive physicochemical changes at variant sites can be good predictors of the null phenotype.

**Figure 1.**
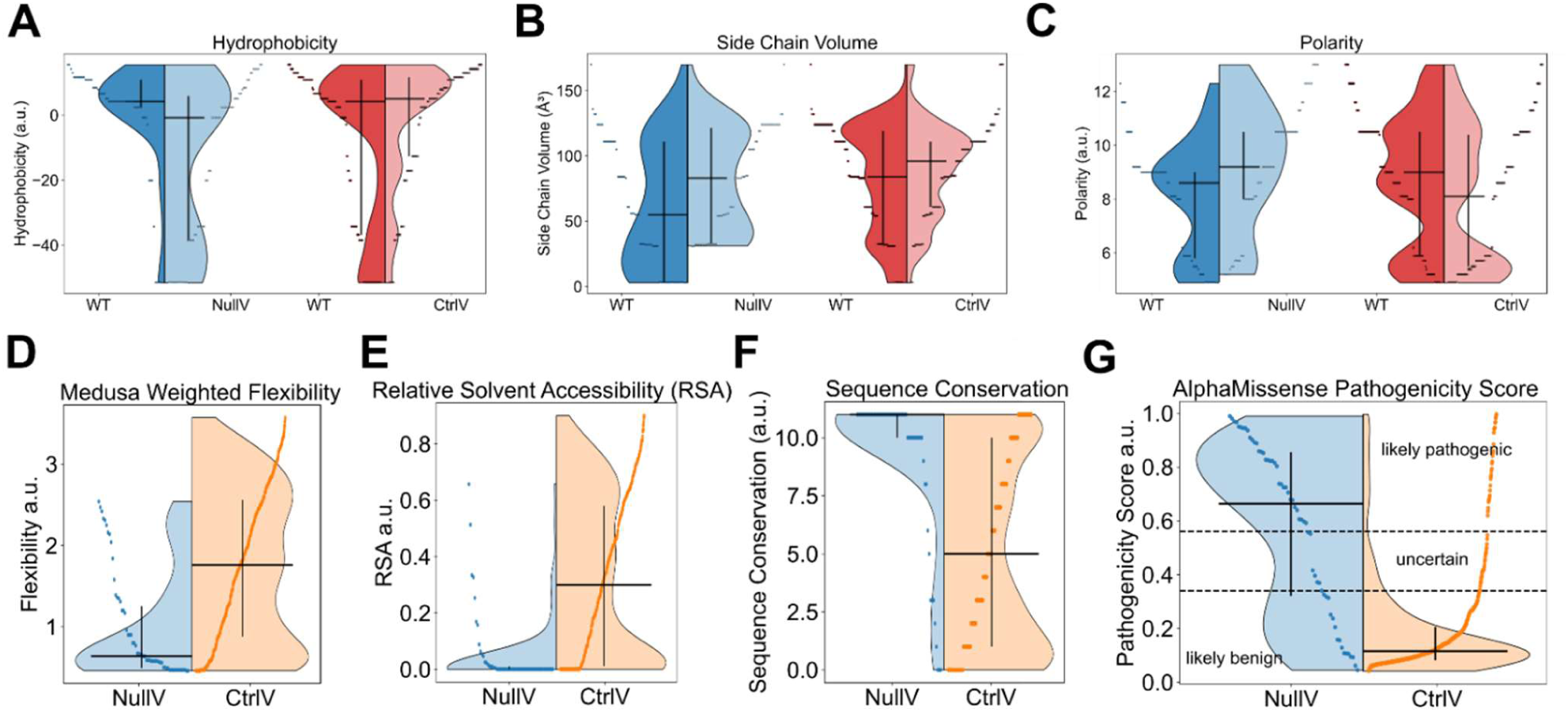
Comparison of structural and mutational parameters between NullV and CtrlV. Each dot represents one amino acid of the respective group; shaded areas represent the density distribution of the data. The horizontal lines indicate medians, and vertical bars indicate the interquartile range. (A) Hydrophobicity distribution of null phenotype variants (NullV, left light blue panel, p<0.001) and control variants (CtrlV, right light red panel, p<0.001) compared to their respective wild-type residues (darker blue and red panels). Lowest values indicate highest hydrophilicity (B) Side chain volume distribution of null phenotype variants (NullV, left panel, p<0.008) and control variants (CtrlV, right panel, p<0.009) compared to their respective wild-type residues. (C) Polarity distribution of null phenotype variants (NullV, left panel, p<0.002) and control variants (CtrlV, right panel, p<0.001) (D) Flexibility (weighted MEDUSA Score; higher value = more flexible), (p < 0.001) (E) Relative solvent accessibility (0 = buried, 1 = fully exposed), (p < 0.001), (F) Sequence conservation score (0 - 11), (p < 0.001), (G) AlphaMissense Pathogenicity score (0-1), (0-0.34 = likely benign, 0.34-0.564 = uncertain, 0.564-1 = likely pathogenic), (p < 0.001). P-values were computed using a two-sided Mann–Whitney U test to assess differences between the compared distributions for each feature. Statistical significance was defined as p < 0.05. A.u. denotes arbitrary units.

We next assessed structural context and sequence conservation. NullV occur preferentially in less flexible regions, with a narrower distribution than CtrlV which are more frequent in semi-flexible and flexible regions (**Figure 1D**). NullV are also enriched at deeply buried, solvent inaccessible sites (**Figure 1E**) and at highly conserved positions (**Figure 1F**), underscoring the sensitivity of structurally constrained residues to mutation. Consistently, AlphaMissense scores classify a substantially higher proportion of NullV as likely pathogenic compared with CtrlV (60.3% of NullV predicted as likely pathogenic, 12.1% uncertain and 27.6% likely benign, and 7.1% of CtrlV predicted as likely pathogenic, 6.0% uncertain and 86.9% likely benign (**Figure 1G**)).

### 3.2 Amino acid composition analysis reveals distinct substitution patterns in null versus control variants

To characterize the physicochemical basis of null (NullV) versus control (CtrlV) single–amino acid variants, we compared their amino acid compositions. Amino acid frequencies were normalized to the complete dataset baseline (set to 1.0), enabling quantitative assessment of residue over- and underrepresentation. Amino acids introduced in null variants (NullV-V) showed marked enrichment of charged residues and depletion of hydrophobic and aromatic residues in comparison to CtrlV-V (**Figure 2A**), consistent with disruption of stabilizing interactions. The amino acid arginine was strongly enriched in NullV-V (4.24× vs 1.04× in CtrlV-V; Δ +3.20), suggesting a prominent role for positive charge in loss of function. The acidic residues aspartate (1.73× vs 0.21×; Δ +1.51) and glutamate (1.71× vs 0.71×; Δ +1.00) were also enriched, indicating accumulation of electrostatic perturbations. Further, proline was enriched in NullV-V (1.72× vs 0.74× in CtrlV-V; Δ +0.99), consistent with destabilising conformational constraints at variant sites. Glycine, leucine, tyrosine and histidine were completely absent from the NullV-V dataset, indicating their introduction is not responsible for any known null phenotype.

**Figure 2.**
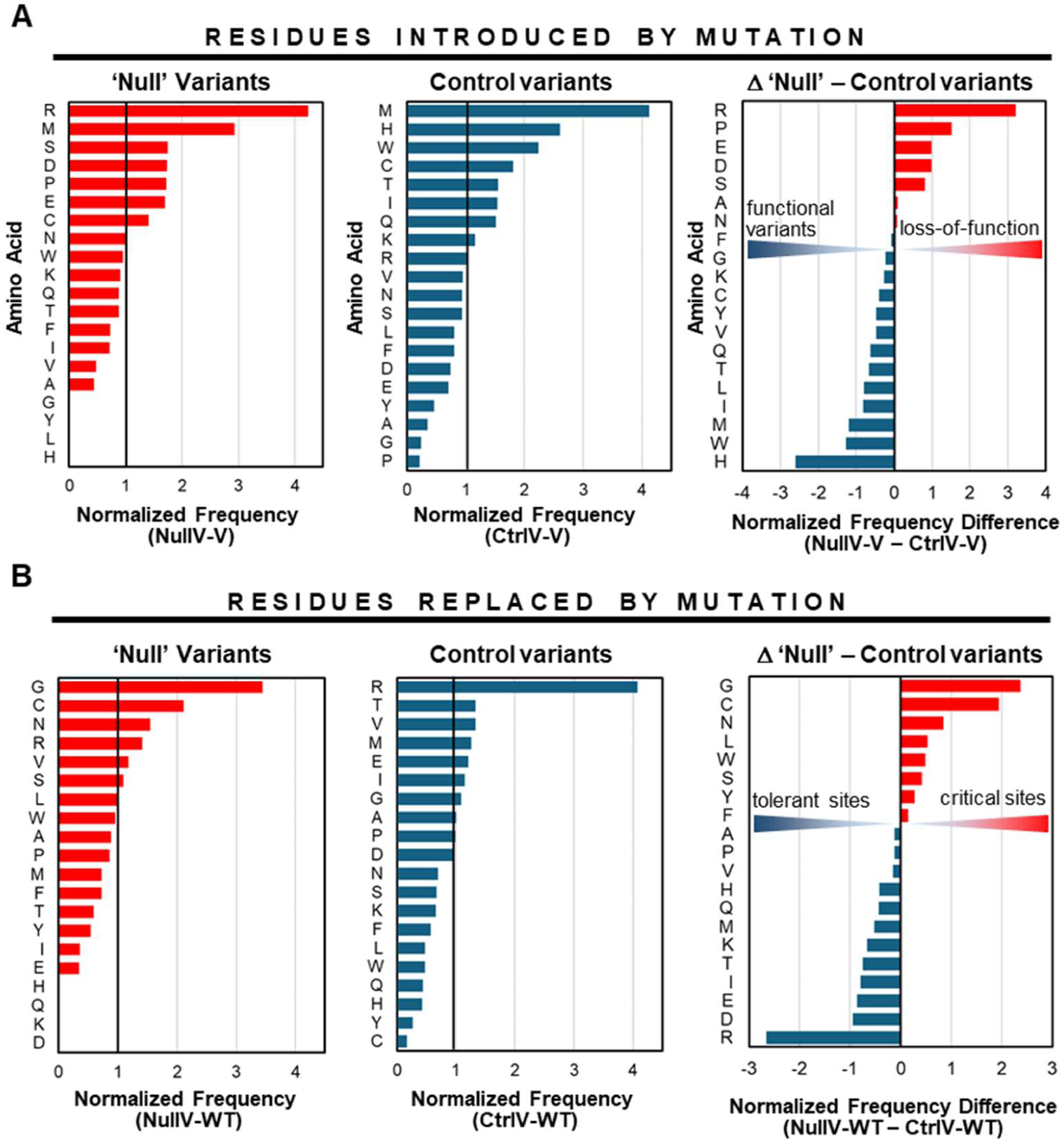
Amino acid frequencies of null variants compared to control variants. All values are normalised to the baseline amino acid frequencies across all full-length protein sequences of the dataset to reveal over- and underrepresentation. N=59 for null variants, and n=481 for control variants. (A) Normalised frequencies of amino acids introduced by null mutations (NullV-V; left panel) compared to control mutations (CtrlV-V; middle panel), and the differences between the two (NullV-V – CtrlV-V; right panel) (B) Normalised frequencies of amino acids replaced by null mutations (NullV-WT; left panel) compared to control mutations (CtrlV-WT; middle panel), and the differences between the two (NullV-WT – CtrlV-WT; right panel).

In contrast, CtrlV-V were enriched in residues associated with stabilizing and aromatic interactions: Methionine showed the highest enrichment (4.13×), while histidine (2.61×) and tryptophan (2.23×) were also increased (0.24×, 0.80×, and 0.47×, respectively), suggesting that tolerated variants retain features supporting flexibility and core packing.

We next analyzed the wild-type residues that get replaced by mutation in each variant class (NullV-WT and CtrlV-WT, **Figure 2B**), to assess whether functional outcome is primarily determined by positional constraints or by substitution chemistry. Null variants preferentially target structurally critical residues, particularly glycine and cysteine (**Figure 2B**). Glycine is strongly enriched (3.45×) in NullV-WT compared with 1.08-fold in CtrlV-WT (Δ +2.37), consistent with disruption of flexible backbone regions. Cysteine-containing positions are also enriched in NullV-WT (2.12× vs 0.17×; Δ +1.94), indicating sensitivity of disulfide-bonded sites to substitution. Also asparagine is overrepresented (1.73× vs 0.74×; Δ +0.99), likely due to the unique role of this amino acid within the restrictive geometry of turns and loops inside proteins^30^. Positions with histidine, glutamine, aspartate, or lysine were completely absent from the NullV-WT group, indicating these amino acids were not replaced in any known null phenotype.

In contrast, control variants preferentially targeted arginine-containing positions (4.07× in CtrlV-WT vs 1.41× in NullV-WT; Δ −2.66), suggesting that arginine’s preferential position at flexible, surface exposed loops makes it inherently more tolerant to amino acid exchanges without any deleterious effect on protein stability.

### 3.3 Prediction of null variants by ML

For classification of null variants (NullV) versus control variants (CtrlV), four ML models were trained using delta hydrophobicity, delta side chain volume, weighted flexibility (MEDUSA), RSA, conservation, and AlphaMissense pathogenicity scores. All models significantly outperformed the dummy baseline (**Figure 3A,B**) in the Dietterich-like analyses, indicating that these descriptors are informative for distinguishing destabilising from neutral variants.

**Figure 3.**
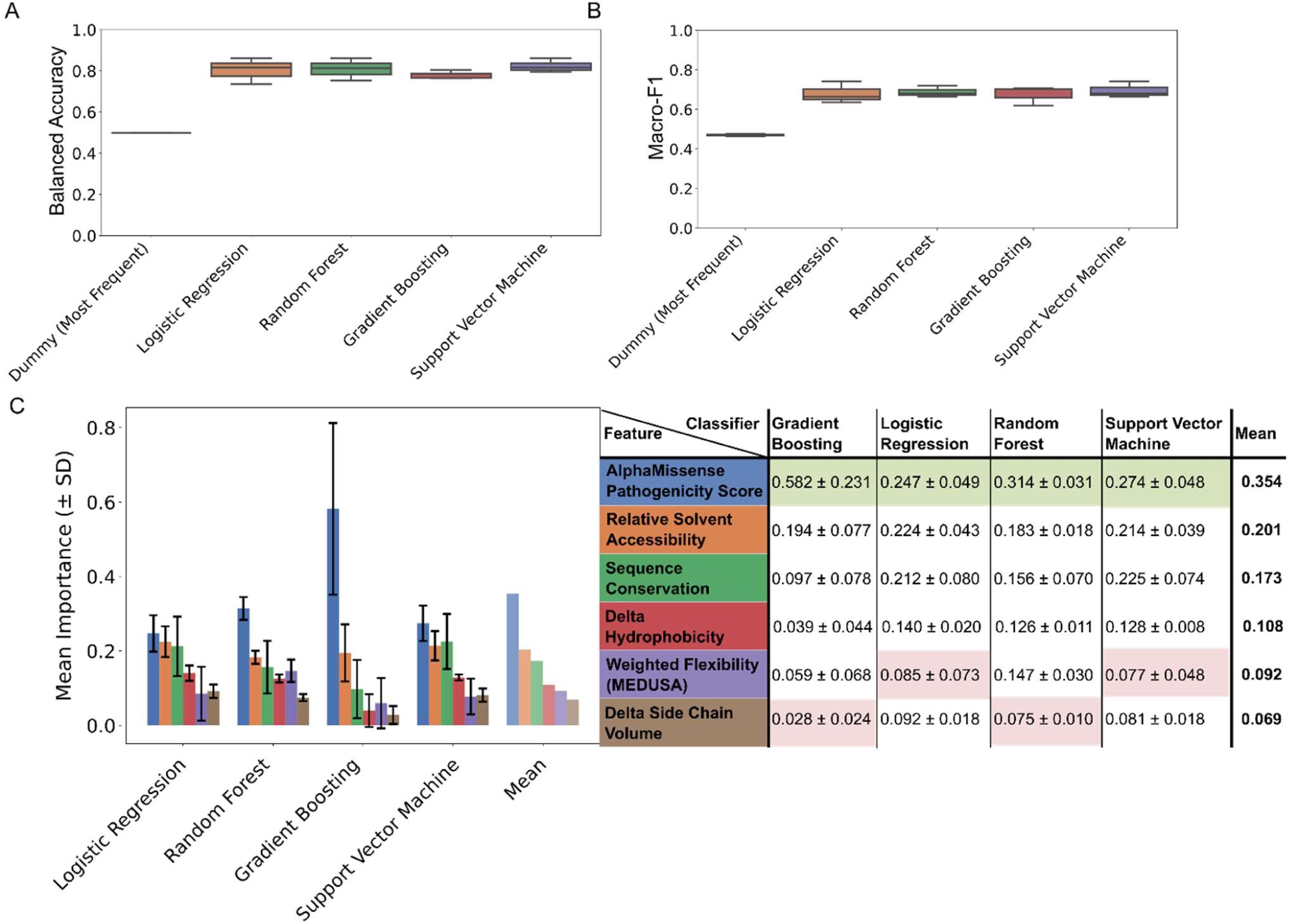
Supervised machine learning model performance and feature importance for NullV versus CtrlV classification. (A) Balanced accuracy (BalAcc) across classifiers. (B) Macro-F1-scores across classifiers. The bar height indicates the mean importance value for a given feature-classifier pair. Error bars correspond to the standard deviation, plotted symmetrically around the mean. A simple baseline classifier that always predicts the most common class in the dataset is included for reference. (C) Feature importance reported as mean ± standard deviation across repeated cross-validation folds (n = 3) for the NullV vs. CtrlV ML classifiers. Highest and lowest values for each individual classifier are highlighted with green and red shading, respectively. Error bars show standard deviation across cross-validation folds, reflecting variability across repeated model training on different subsets of the data.

Among the supervised classifiers tested, SVM achieved the highest median performance (BalAcc 0.82; Macro-F1 0.70) followed closely by Random Forest (BalAcc 0.81; Macro-F1 0.69), and Logistic Regression (BalAcc 0.80, Macro-F1 0.68). Gradient Boosting (BalAcc 0.78, Macro-F1 0.68) performed slightly worse. No statistically significant pairwise differences were detected among the four classifiers after correction. These results demonstrate that both ensemble methods and SVM are well suited to separate NullV from CtrlV variants, with SVM showing the best performance.

For BalAcc and Macro-F1, supervised classifiers outperformed the dummy baseline in the Dietterich-like analyses, whereas the Alpaydin-like tests did not consistently confirm significance against the dummy baseline. Feature importance analysis identified the AlphaMissense pathogenicity score as the dominant predictor (mean feature importance [FI] 0.354) ranking highest all four models (**Figure 3C**). Relative solvent accessibility (mean FI 0.204) and conservation (mean FI 0.173) were the next most informative features (**Figure 3C**). In contrast, MEDUSA weighted flexibility (mean FI 0.092), and the biophysical change features such as hydrophobicity difference (‘delta’, mean FI 0.108), and side chain volume difference (mean FI 0.069) contributed less strongly. Nonetheless, these features distinguish NullV and CtrlV at distribution level (**Figure 1**), suggesting they provide complementary information to structure- and conservation-based predictors.

As AlphaMissense pathogenicity score was the most influential feature, we performed an ablation analysis excluding it. Model performance decreased only moderately, indicating that predictive performance did not depend primarily on this feature alone (see **Supplemental data** and **Supplementary Table 2**.)

### 3.4 Comparison of amino acid determinants in antigenic and control variants

#### 3.4.1 Qualitative Analysis

We next compared physicochemical and structural properties of amino acids in antigenic variants with control variants lacking known antigenicity (CtrlV) (**Figure 4**).

**Figure 4.**
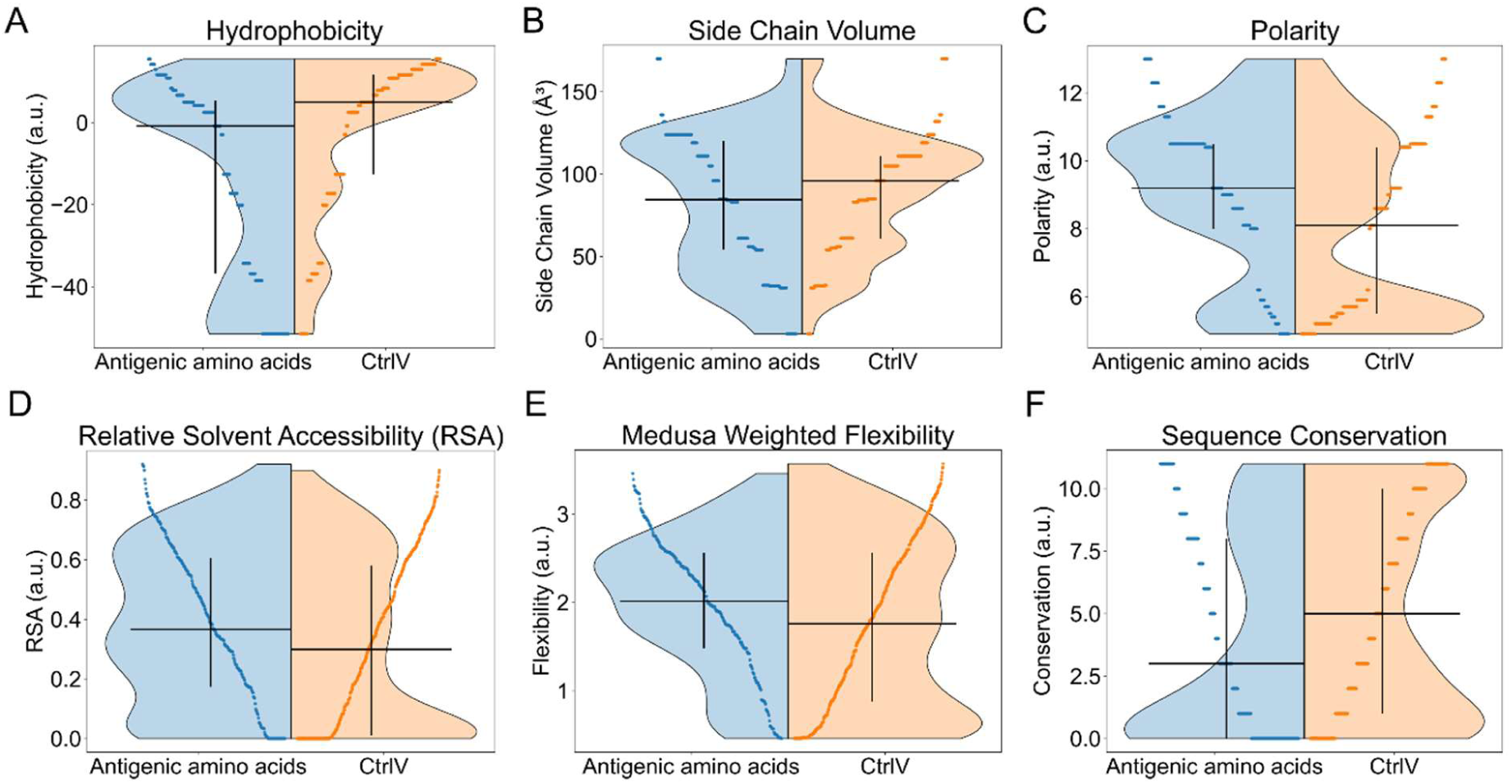
Distribution of structural and biophysical properties across blood group antigens and control variants. Violin plots showing the distributions of (A) hydrophobicity (p<0.001), (B) side chain volume (p=0.129), (C) polarity (p<0.001), (D) relative solvent accessibility (p<0.001), (E) predicted flexibility (p=0.002), and (F) sequence conservation (p<0.001), for antigenic and control amino acids across all included blood group systems. Values are displayed on the left for antigenic residues (AgWT and AgV, blue) and on the right for non-antigenic residues (CtrlV, orange). The blue vertical bar left and right of each violin plot denotes the interquartile range, with the horizontal line indicating the median. Data points are overlaid to illustrate sample density. P-values were computed using a two-sided Mann–Whitney U test to assess differences between antigenic amino acids and CtrlV distributions for each feature. Statistical significance was defined as p < 0.05. A.u. denotes arbitrary units.

Antigenic variants showed lower hydrophobicity, increased polarity, and moderately smaller side chain volumes compared with CtrlV (**Figure 4A-C**), consistent with an enrichment of solvent-compatible residues.

Both the median predicted flexibility (2.02 vs 1.76) and the median relative solvent accessibility (0.37 vs 0.30) were higher in antigenic residues. 22.3% of antigenic residues and 37.2% of control residues were completely buried (RSA < 0.15), and 65.0% of antigenic residues and 53.8% of control residues were exposed (RSA > 0.25)(**Figure 4D,E**). This might reflect the fact that many antigens represent amino acid side chains on or near the surface of protein molecules where they contribute directly or indirectly to solvent-exposed epitopes. The median conservation level of antigenic residues was lower than those of control residues (**Figure 4F**), indicating that antigenic sites reside in evolutionarily more variable regions of proteins than the control variants.

#### 3.4.2 Amino acid composition distinguishes antigenic from non-antigenic sites

We next compared amino acid composition between antigenic residues (AgWT and AgV) and non-antigenic control variants (CtrlV) to identify features associated with immunogenic epitopes (**Figure 5**). Frequencies were again normalized to a proteome-wide baseline (1.0).

**Figure 5.**
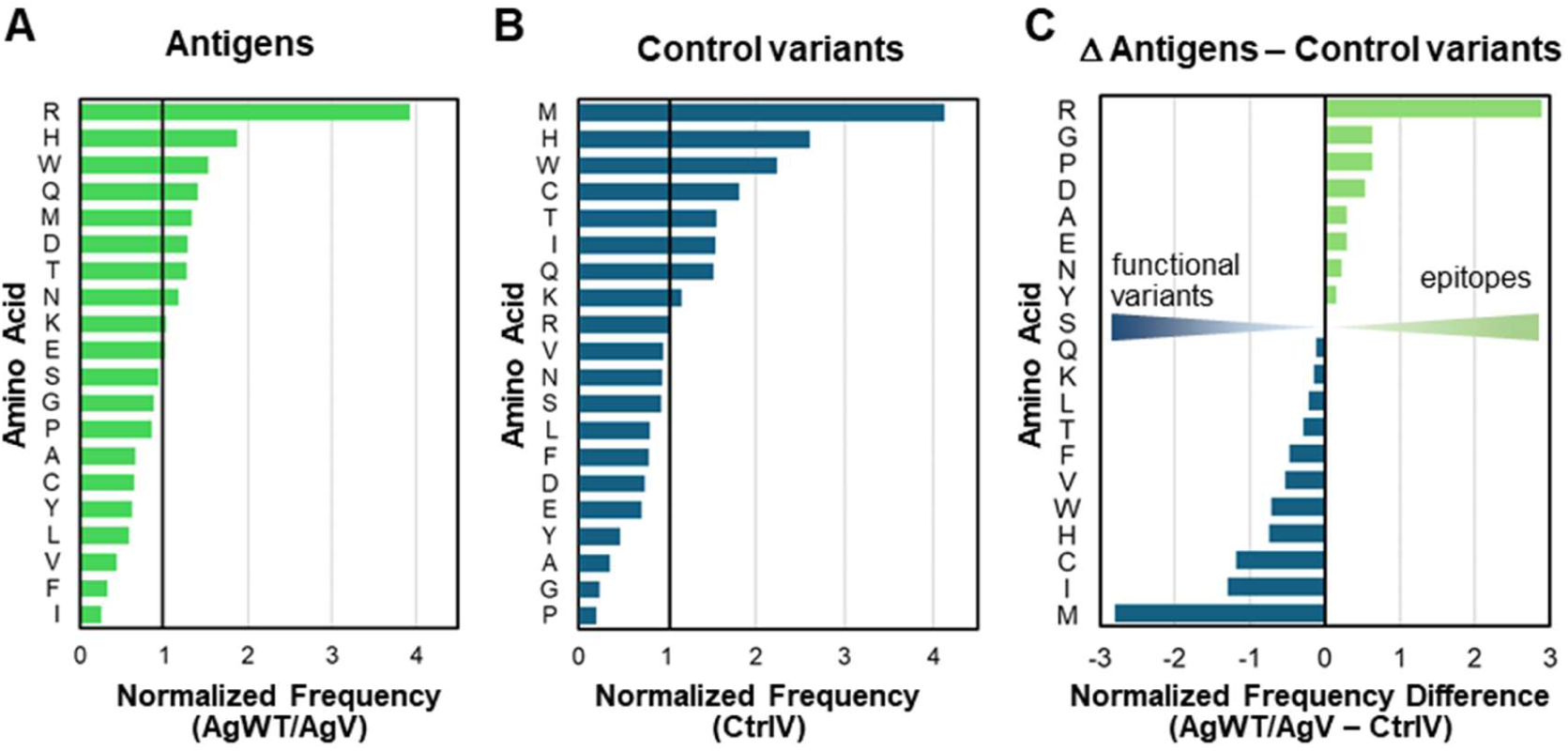
Amino acid frequencies of antigenic variants compared to control variants. All values are normalised to the baseline amino acid frequencies across all full-length protein sequences of the dataset to reveal over- and underrepresentation. N=260 for antigenic variants (comprising AgWT and AgV), and n=481 for control variants. (A). Normalised frequencies of antigen-defining amino acids (AgWT/AgV). (B) Normalised frequencies of control variants (CtrlV) (C) Differences between normalised frequencies of antigens and control amino acids (AgWT/AgV - CtrlV).

Antigenic residues showed a marked overrepresentation of arginine (3.93×) compared with controls (1.04×; Δ +2.89), pointing out the positively charged arginine side chain as a potent mediator of antigen/epitope recognition. Glycine and proline – although overall underrepresented in both antigens and controls (**Figure 5A,B**), were more common in antigens than in controls, consistent with altered flexibility or structural perturbation at antigenic sites. In contrast, hydrophobic and aromatic residues were depleted in antigens but enriched in controls. Methionine was strongly enriched in controls (4.13× vs 1.33× in antigens; Δ −2.80), while isoleucine (1.54× vs 0.24×; Δ −1.30) and cysteine (1.82× vs 0.64×; Δ −1.18) showed similar trends, indicating depletion of core-stabilizing residues at antigenic positions. Additional hydrophobic residues, including leucine, valine, and phenylalanine, were likewise underrepresented in antigens, reflecting the fact that they rarely appear on protein surfaces.

#### 3.4.3 Prediction of antigenicity by ML

For the classification of antigenic residues versus control variants, supervised models were trained using hydrophobicity, conservation, weighted flexibility (MEDUSA), relative solvent accessibility and side chain volume data. All classifiers significantly outperformed the baseline (**Figure 6**), indicating that these features capture signal relevant to antigenicity. Logistic Regression, Random Forest and SVM achieved a slightly higher performance (BalAcc ∼0.63,) than Gradient Boosting (BalAcc ∼0.60). Overall performance was reduced compared with the NullV vs. CtrlV classification, indicating greater difficulty in distinguishing antigenic variants from neutral variants based on the features provided. As for the null classification, no statistically significant pairwise differences were detected among the four classifiers after correction.

**Figure 6:**
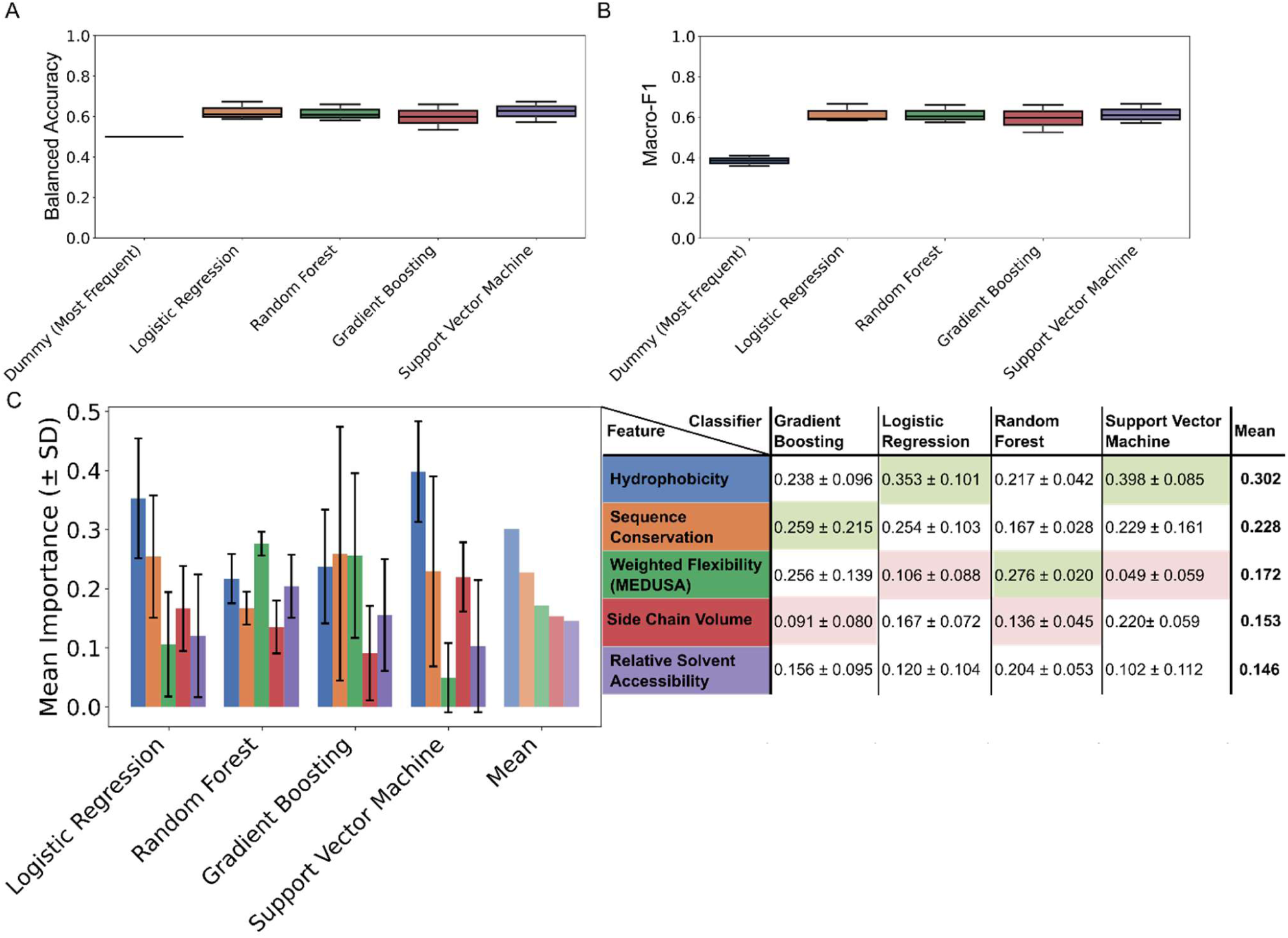
Model performance for antigenic vs. non-antigenic classification. (A) Box plots of BalAcc across classifiers. (B) Box plots of Macro-F1-scores across classifiers for antigenic vs. non-antigenic classification. The bar height indicates the mean importance value for a given feature-classifier pair. Error bars correspond to the standard deviation, plotted symmetrically around the mean. A simple baseline classifier that always predicts the most common class in the dataset is included for reference. (C) Feature importance as mean ± standard deviation across repeated cross-validation folds (n = 3) for the antigenic vs non-antigenic classifiers.

Feature importance analysis identified hydrophobicity (mean FI 0.302), conservation (0.228) and flexibility (0.172) as the most influential predictors (**Figure 6C**). This is partially in line with the finding that conservation showed the largest difference between median values for antigenic and non-antigenic variants (**Figure 4F**), supporting its role as a key discriminative feature. Side chain volume contributed less strongly, consistent with the NullV vs. CtrlV analysis. Notably, hydrophobicity ranked highest in the Logistic Regression and SVM classifiers, indicating that it contributed strongly to discrimination in the linear models. In contrast, conservation and flexibility were more prominent in tree-based models (**Figure 6C**) indicating that the relative importance of individual descriptors depended on the classifier used. Variability in feature importance rankings across models suggests no single uniform importance pattern across classifiers.

## 4 Discussion

This study provides the first cross-system, residue-level comparison of structural, biophysical, and evolutionary features associated with antigenic and null phenotypes. Extending observations from individual systems, we find that loss of antigen expression and antigenicity are governed by distinct and consistent molecular constraints.

Variants leading to null phenotypes preferentially occur at highly conserved, buried hydrophobic sites with low flexibility and solvent accessibility. Disruption of such sites, particularly those involving glycine and cysteine residues, likely interferes with backbone flexibility or disulfide bond formation leading to pronounced structural destabilisation. The observed decrease in hydrophobicity and depletion of aromatic residues in null variants further support disruption of the hydrophobic core. The high frequency of mutations to arginine in null variants highlights electrostatic perturbation as a key contributor to loss of antigen expression.

In contrast, control variants that preserve expression tend to occur at more permissive positions including highly flexible and fully solvent-exposed regions and maintain features compatible with structural stability. Together, these findings indicate that loss of antigen expression reflects the interplay between positional context and substitution chemistry, with structurally critical sites being particularly sensitive to disruptive changes.

Overall, antigenic variants differed more subtly from control variants than null variants, suggesting that antigenicity is governed by more complex and context-dependent mechanisms. Compared with controls, antigenic amino acids tend to be less hydrophobic, more polar and reside in more solvent accessible and more flexible regions. They are enriched in arginine, suggesting that electrostatic properties contribute to epitope formation. Localized charge might shape antigenic properties, potentially through modulation of protein–antibody interactions. The scarcity of hydrophobic and aromatic residues amongst antigens further supports exclusion from structurally constrained core regions and instead a requirement for surface exposure (or vicinity) and conformational adaptability in antigen formation. Notably, antigenic residues are less conserved than control variants, a fact that cannot be explained by structural tolerance to mutation alone. In fact, antigens may evolve under selective pressures related to host–pathogen or immune interactions. This is in line with the concept of balancing selection maintaining diversity at exposed protein regions.

Encouragingly, our four models could discriminate both null and antigenic variants from benign, non-antigenic variants from a limited but highly evidence-based training dataset of all known single amino acid substitutions reported by the ISBT. With a BalAcc of up to 0.82 for null phenotype prediction and up to 0.63 for antigenicity prediction, the performance of the predictions is, however, inferior to genome-based blood typing^6^. Classification of null variants was driven primarily by pathogenicity scores, solvent accessibility, and conservation, consistent with structural destabilization. Similar patterns of feature importance across all 24 blood group systems suggest that the determinants of antigen loss are broadly generalizable across proteins. In contrast, antigenicity prediction relied more diffusely on hydrophobicity, conservation, and flexibility, with less consistent feature importance across models. This difference is reflected in lower predictive performance compared to null variant prediction, indicating that antigenicity arises from a combination of physicochemical properties and structural context rather than a single dominant factor. This is consistent with established models of epitope formation, in which surface physicochemical properties and local conformational dynamics jointly modulate antigen exposure and antibody binding potential^31^. Additional features such as detailed surface geometry, dynamic conformational changes, or explicit epitope descriptors, may be required to improve accuracy. The relatively low contribution of solvent accessibility in the current models seems in contrast to a recent finding that the four most immunogenic blood group antigens arise from substitutions at significantly more buried protein regions than five less immunogenic antigens^17^. This may be due to our binary (yes/no) classification, which does not account for varying degrees of antigenicity.

Several limitations should be considered. Despite its diversity across blood groups, the dataset remains biased toward experimentally well-characterized systems such as Rh and Kell. Systems like Lutheran are quantitatively well represented but lack comparable experimental validation. In order to capture dynamic conformational changes induced by single mutations that may influence protein stability or epitope exposure *in vivo*, molecular dynamics simulations (MD) and (folding) free-energy minimization calculations would provide valuable additional determinants^32^. Incorporating a quantitative measure of immunogenicity into the training set could allow for a more nuanced prediction of the expected medical impact of novel antigens. However, immunogenicity is inherently difficult to quantify due to its dependence on a complex interplay of host-specific factors, antigen properties, and contextual variables, which vary widely and resist reduction to a single standardized metric. Emerging AI-driven epitope prediction models^33^ offer further future options to refine automatic prediction.

In summary, single-amino-acid substitutions underlying blood group phenotypes follow consistent structural and biophysical principles across systems. Null phenotypes arise from substitutions at structurally constrained and evolutionarily conserved sites that disrupt protein stability, whereas antigenic variants are enriched at more flexible, solvent-accessible, and evolutionarily variable positions. Although antigenicity is inherently more difficult to predict than loss of expression, the consistent importance of hydrophobicity, conservation, and structural context supports the development of interpretable, structure-aware approaches for evaluating novel missense variants in transfusion medicine.

## Supporting information

Fasta Sequence Alignments

## Acknowledgments

We are very grateful to Dr Gabriele Mayr (IKMB Kiel, Germany) for critical reading of the manuscript and numerous valuable comments. The Institute of Translational Medicine at the Private University in the Principality of Liechtenstein is supported by the Hans Groeber-Stiftung (Vaduz, Principality of Liechtenstein); and the Tarom Foundation (Schaan, Principality of Liechtenstein).

## 5 Authorship Contributions

Conceptualization, supervision: (MB,CG, JS). Data collection, data analysis, investigation, validation: (MB,ACK,JS). Visualization: (ACK). Writing – original draft: (ACK). Writing – review & editing (MB, JS, CG). Made substantial, direct and intellectual contributions to the work, and approved it for publication: (ACK,MB,JS,CG).

## 6 Conflict of Interest

CG acts as a consultant to Inno-Train GmbH, Kronberg im Taunus, Germany, a provider of genotyping kits for molecular blood group diagnostics since 1998. CG holds the European and US patents P3545102 and US20190316189 on the “‘Determination of the genotype underlying the S-s-U-phenotype of the MNSs blood group system”’. The other authors declare that the research was conducted in the absence of any commercial or financial relationships that could be construed as a potential conflict of interest.

## Notes

https://github.com/ackranz/blood-phenotype-ml-supplement

