## Supplemental Material PDF for "Determinants of Blood Group Antigen Expression and Prediction of Phenotypes by Machine Learning"

**Contents**
**Supplemental Methods**
**Supplemental Tables 1-3**
**Supplemental Analyses**
- *Multicollinearity Analysis*
- *Ablation Analysis without AlphaMissense*

**<u>Additional Supplemental files:</u>**

| | |
|---|---|
| **ML-supplement.zip** | ML scripts and documentation |
| **Supplemental_Tables.xls** | Extended Supplemental Table1 and Supplemental Table 2 |
| **Alignment_files_fasta.zip** | Alignments used for sequence conservation analysis |

### Supplemental Methods

All analyses were performed in Python 3.12 using numpy [2.0.2], pandas [2.2.3], scipy [1.14.1], scikit-learn [1.5.2], matplotlib [3.9.2], and imbalanced-learn [0.14.0]. The complete analysis code is available as Supplemental file **ML-Supplement.zip** as well as in the GitHub repository https://github.com/ackranz/blood-phenotype-ml-supplement, with scripts and documentation sufficient to reproduce the workflows reported.

### Supplemental Tables

Extended supplemental tables are available for download as **Supplemental_Tables.xls**

**Supplemental** Table 1: Overview of structural models used in this study.

| ISBT System # | Blood group | Symbol | Gene | UniProt | Structural model source | Chain(s) | Notes |
|---|---|---|---|---|---|---|---|
| 2 | MNS | MNS | GYPA | P02724 | AlphaFold Database[1] | A | |
| 2 | MNS | MNS | GYPB | P06028 | AlphaFold Database[1] | A | Truncated to residues 20-91 |

| | | | | | | | |
|---|---|---|---|---|---|---|---|
| 4 | RhD | RHD | RHD | Q02161 | AlphaFold Database[1] | A,B,C | |
| 5 | Lutheran | LU | BCAM | P50895 | AlphaFold Database[1] | A | |
| 6 | Kell | KEL | KEL | P23276 | AlphaFold Server[2] | A | |
| 8 | Fy (Duffy) | FY | ACKR1 | Q16570 | AlphaFold Database[1] | A | |
| 9 | Kidd | JK | SLC14A1 | Q13336 | PDB 6qd5 | A,B,C | Trimer |
| 10 | Diego | DI | SLC4A1 | P02730 | PDB 7uz3, Alphafold Database[1] | A,B,C,D | TM domains were taken from PDB 7uz3, and soluble domains from Alphafold. |
| 11 | Yt | YT | ACHE | P22303 | PDB 4ey4 | A,B | Dimer |
| 13 | Scianna | SC | ERMAP | Q96PL5 | AlphaFold Database[1] | A | |
| 14 | Dombrock | DO | ART4 | Q93070 | AlphaFold Database[1] | A | |
| 15 | Colton | CO | AQP1 | P29972 | PDB 7uze | A,B,C,D | Tetramer |
| 16 | Landsteiner -Wiener (LW) | LW | ICAM4 | Q14773 | AlphaFold Database[1] | A | Model was truncated to residues 23-271 |
| 20 | Gerbich (Ge) | GE | GYPC | P04921 | AlphaFold Database[1] | A | |
| 21 | Cromer | CROM | CD55 | P08174 | PDB 1ojv | A, B | Dimer |
| 23 | Indian | IN | CD44 | P16070 | AlphaFold Server[2] | A | Predicted structure of relevant isoform 12 |
| 24 | Ok | OK | BSG | P35613 | AlphaFold Database[1] | B | |
| 25 | Raph | RAPH | CD151 | P48509 | AlphaFold Database[1] | A | |
| 26 | John Milton Hagen | JMH | SEMA7A | O75326 | PDB 3nvq | A,E | Heterotetramer with 2 copies of SEMA7A and 2 copies of Plexin-C1 |
| 30 | Rh-associated glycoprotein | RHAG | RHAG | Q02094 | PDB 7uzq | B,C | Heterotrimer of $RHAG_2$-$RHCE_1$, bound to Ankyrin |
| 35 | CD59 | CD59 | CD59 | P13987 | PDB 2j8b | A | |
| 36 | Augustine | AUG | SLC29A1 | Q99808 | PDB 6ob7 | A | |
| 37 | Kanno | KANNO | PRNP | P04156 | PDB 4klm | A | |
| 39 | CTL2 | CTL2 | SLC44A2 | Q8IWA5 | AlphaFold Database[1] | A | CTL2-P1 isoform (Isoform 3 - Q8IWA5-3 in UniProt) |

[1]AlphaFold Protein Structure Database: https://alphafold.ebi.ac.uk/

[2]AlphaFoldServer: https://alphafoldserver.com/

This table is also available in the file **Supplemental_Data.xls** including the reference amino acid sequences.

**Supplemental Table 2: Full dataset file**

Supplemental Table 2 comprises the full dataset of all variants and prameters used in the analysis. It is available in the file **Supplemental_Data.xls**

**Supplemental Table 3: Balanced accuracy and Macro F1 scores for the tested classifiers with and without AlphaMissense pathogenicity score feature**

| Classifier | Balanced accuracy without AlphaMissense | Balanced accuracy with all features | Macro F1 without AlphaMissense | Macro F1 with all features |
|---|---|---|---|---|
| **Dummy** | 0.5 | 0.5 | 0.471 | 0.471 |
| **Gradient Boosting** | 0.731 | 0.778 | 0.651 | 0.676 |
| **Logistic Regression** | 0.794 | 0.804 | 0.651 | 0.681 |
| **Random Forest** | 0.788 | 0.809 | 0.671 | 0.689 |
| **SVM** | 0.799 | 0.823 | 0.649 | 0.696 |

### Supplemental Analyses

*Multicollinearity analysis*

We assessed multicollinearity using pairwise Pearson correlations and variance inflation factors (VIFs). For the predictor CtrlV vs NullV we used the following features: Delta hydrophobicity, delta side chain volume, relative solvent accessibility, MEDUSA weighted flexibility class, conservation, predicted pathogenicity of mutation. For the predictor CtrlV vs Ag we used the features: Hydrophobicity, side chain volume, relative solvent accessibility, conservation, MEDUSA weighted flexibility class.
The VIF values were low (maximum VIF 2.10 for CtrlV vs NullV and 1.78 for CtrlV vs Ag), indicating no problematic multicollinearity. The strongest pairwise correlation in both tasks was between relative solvent accessibility and MEDUSA weighted flexibility class (r = 0.702 and r = 0.646, respectively).

An earlier version included delta polarity (CtrlV vs NullV), which increased the maximum VIF to 6,01 (CtrlV vs NullV), with the strongest correlation observed between delta hydrophobicity and delta polarity (r = 0.743). Adding polarity (CtrlV vs Ag) increased the maximum VIF to 9.34, and Hydrophobicity and Polarity were strongly correlated (r = 0.824). Therefore, polarity and delta polarity were not included in the final feature set.

*Ablation analysis without AlphaMissense*

For the prediction of null variants AlphaMissense pathogenicity scores is the most influential feature across classifiers. To assess whether predictive performance depends primarily on the AlphaMissense score, we performed an ablation analysis in which the models were re-trained without AlphaMissense and re-evaluated using the same grouped repeated nested cross-validation and paired significance-testing framework against the dummy baseline. The results without the AlphaMissense decrease in balanced accuracy and F1-Macro scores, but not to an extent which would lead to the conclusion that predictive performance mainly depends on the AlphaMissense pathogenicity score feature. With all features present (delta hydrophobicity, delta sidechain volume, solvent accessibility, weighted flexibility, conservation, AlphaMissense pathogenicity score) the balanced accuracy for the four tested classifiers lies between 0.778 and 0.823 (Macro F1 0.676 to 0.696), whereas leaving out AlphaMissense pathogenicity scores it drops to between 0.731 and 0.799 (Macro F1 0.649 to 0.671) (see **Supplemental Table 3**).
