## Supplementary material for "Determinants of Blood Group Antigen Expression and Prediction of Phenotypes by Machine Learning": ML Supplement: feature_importance_visualization.py

```
import pandas as pd
import numpy as np
import matplotlib.pyplot as plt

# === 1) Load your data ===
# Change this path if your file lives elsewhere
#csv_path = "train_delta_features_ctrlv_nullv_results_undersampling_LR_04_grouped_feat_imp_per_fold/feature_importances_variance_summary.csv"
csv_path = "train_delta_features_{TASK.lower()}}_results_undersampling_LR_04_grouped_01_set69_v01/feature_importances_variance_summary.csv"#train_classifiers_agv_agwt_ctrlv_resultsperfold_joinedfile_KGGGIDADSRRLKLOCR_26__18.11/feature_importances_variance_summary.csv"
df = pd.read_csv(csv_path)

# Ensure we have a standard deviation column; compute it from variance if needed
if "std" not in df.columns and "var" in df.columns:
    df["std"] = np.sqrt(df["var"])
elif "std" not in df.columns:
    raise ValueError("CSV must contain either 'std' or 'var' to draw error bars.")

# === 2) Order categories (optional but helps consistent grouping) ===
classifiers = df["Classifier"].unique()
features = df["Feature"].unique()

# === 3) Build a grouped bar plot with error bars ===
x = np.arange(len(features))                 # feature positions
width = 0.8 / len(classifiers)               # bar width for each classifier

plt.figure(figsize=(12, 6))

for i, clf in enumerate(classifiers):
    subset = df[df["Classifier"] == clf]
    # Align bars with feature order
    subset = subset.set_index("Feature").reindex(features).reset_index()

    plt.bar(
        x + i * width,
        subset["mean"].values,
        yerr=subset["std"].values,
        width=width,
        label=clf,
        capsize=5
    )

plt.title("Mean Feature Importance per Classifier")
plt.xlabel("Feature")
plt.ylabel("Mean Importance (± SD)")
plt.xticks(x + width * (len(classifiers) - 1) / 2, features, rotation=45, ha="right")
plt.legend(title="Classifier")
plt.tight_layout()

# === 4) Save (and/or show) ===
plt.savefig("train_delta_features_ctrlv_nullv_results_undersampling_LR_04_grouped_01_set62_v07/feature_importances_grouped_errorbars.png",  dpi=300)
            #"train_delta_features_ctrlv_nullv_results_undersampling_LR_04_grouped_feat_imp_per_fold/feature_importance_grouped_errorbars.png", dpi=300)
# plt.show()
```
