## Supplementary material for "Determinants of Blood Group Antigen Expression and Prediction of Phenotypes by Machine Learning": ML Supplement: README.md

# Blood Phenotype ML Supplement

This repository contains supplementary Python scripts for training, evaluating, and visualising machine-learning classifiers for blood phenotype prediction.

The workflow uses group-aware nested cross-validation, feature-importance analysis, statistical comparison of classifiers, and post-hoc visualisations.

## Contents

| File | Description |
|---|---|
| `train_classifier_12_05_26.py` | Main training script. Runs grouped repeated nested cross-validation, trains multiple classifiers, saves metrics, confusion matrices, feature importances, SHAP plots, and statistical test outputs. |
| `boxplot_visualisations_with_stat_sign_a.py` | Creates boxplots and bar charts from `fold_results.csv`, including optional significance annotations if pairwise significance results are available. |
| `feature_importance_visualization.py` | Creates grouped feature-importance visualisations from `feature_importances_variance_summary.csv`. |
| `requirements.txt` | Python package requirements needed to run the scripts. |

## Main analysis

The main script trains and evaluates the following classifiers:

- Dummy baseline
- Logistic Regression
- Random Forest
- Gradient Boosting
- Support Vector Machine

The current configuration runs the `CTRLV_VS_NULLV` task and uses a grouped repeated cross-validation design with 5 repeats, 3 outer folds, and 2 inner folds.

Input data are expected in:

```text
merged_variant_data_78_a.csv
