## Supplementary material for "Determinants of Blood Group Antigen Expression and Prediction of Phenotypes by Machine Learning": ML Supplement: boxplot_visualisations_27.08.py

```
import os
import numpy as np
import pandas as pd
import matplotlib.pyplot as plt
import seaborn as sns

# =========================
# Config
# =========================
# Path to your per-fold results CSV
file_path = "train_delta_features_ctrlv_nullv_results_undersampling_LR_04_grouped_feat_imp_per_fold/fold_results.csv"
    #"train_classifiers_agv_agwt_ctrlv_resultsperfold_joinedfile_KGGGIDADSRRLKLOCR_26_selected_features_grouped/fold_results_agv_agwt_ctrlv.csv"#"train_delta_features_ctrlv_nullv_results_untersampling/fold_results.csv"#"train_classifiers_agv_agwt_ctrlv_resultsperfold_joinedfile_KGGGIDADSRRLKLOCR_26_selected_features_c/fold_results_agv_agwt_ctrlv.csv"

# Output directory = same as the CSV's directory
out_dir = os.path.dirname(file_path)
os.makedirs(out_dir, exist_ok=True)

# Matplotlib font size
plt.rcParams.update({'font.size': 14})

# =========================
# Load
# =========================
df = pd.read_csv(file_path)

# =========================
# Choose metrics dynamically (handles old & new CSVs)
# =========================
all_candidate_metrics = [
    "Train Accuracy", "Test Accuracy", "Test Precision", "Test Recall", "Test F1-Score",
    "F1-Macro on test data", "Balanced Accuracy on test data"  # NEW metrics
]
metrics = [m for m in all_candidate_metrics if m in df.columns]

if not metrics:
    raise ValueError("No known metric columns found in the CSV. "
                     "Expected one of: " + ", ".join(all_candidate_metrics))

# =========================
# Boxplots per metric
# =========================
for metric in metrics:
    plt.figure(figsize=(10, 6))
    sns.boxplot(x="Classifier", y=metric, data=df, hue="Classifier", palette="Set3")
    plt.title(f"{metric}")
    plt.ylabel(metric)
    plt.xlabel("Classifier")
    plt.xticks(rotation=45, ha="right")
    plt.ylim(0, 1)
    plt.legend([], [], frameon=False)  # hide legend (duplicate of x labels)
    plt.subplots_adjust(bottom=0.35)   # more space for rotated labels

    plot_file_path = os.path.join(out_dir, f"{metric.replace(' ', '_')}_boxplot.png")
    plt.savefig(plot_file_path, dpi=200, bbox_inches="tight")
    plt.close()

print("Box plots have been saved successfully.")

# =========================
# Summary table: mean & std per metric/classifier
# =========================
if metrics:
    agg_spec = {m: ['mean', 'std'] for m in metrics}
    summary_data = df.groupby('Classifier').agg(agg_spec).reset_index()

    # Flatten MultiIndex columns: ("Test Accuracy","mean") -> "Test Accuracy Mean"
    new_cols = []
    for c in summary_data.columns:
        if isinstance(c, tuple):
            if c[1] == '':
                new_cols.append(c[0])  # e.g. "Classifier"
            else:
                new_cols.append(f"{c[0]} {'Mean' if c[1] == 'mean' else 'Std Dev'}")
        else:
            new_cols.append(c)
    summary_data.columns = new_cols

else:
    raise ValueError("No known metric columns found in the CSV. ")

# Save summary table
summary_csv_path = os.path.join(out_dir, "metrics_summary_by_classifier.csv")
summary_data.to_csv(summary_csv_path, index=False)

# =========================
# Bar charts (mean ± std) per metric
# =========================
classifiers = summary_data['Classifier']
plt.rcParams.update({'font.size': 22})

for metric in metrics:
    mean_col = f"{metric} Mean"
    std_col  = f"{metric} Std Dev"

    if mean_col not in summary_data.columns:
        continue

    means = summary_data[mean_col]
    std_devs = summary_data[std_col] if std_col in summary_data.columns else None

    x_pos = np.arange(len(classifiers))

    plt.figure(figsize=(12, 8))
    plt.bar(x_pos, means, yerr=std_devs, align='center', alpha=0.7, ecolor='black', capsize=11)
    plt.xticks(x_pos, classifiers, rotation=45, ha='right')
    plt.xlabel('Classifier')
    plt.ylabel(metric)
    plt.title(f'{metric} Across Classifiers')
    plt.ylim(0, 1)
    plt.tight_layout()

    plot_file_path = os.path.join(out_dir, f"{metric.replace(' ', '_')}_bar_chart.png")
    plt.savefig(plot_file_path, dpi=200, bbox_inches="tight")
    plt.close()

print("Bar charts have been saved successfully.")

# =========================
# OPTIONAL: Confusion matrix heatmaps (if CSVs exist)
# Expects files named like: confusion_matrix_<ClassifierNameWithUnderscores>.csv
# created by your training script per classifier.
# =========================
unique_classifiers = df['Classifier'].unique()
for clf in unique_classifiers:
    cm_file = os.path.join(out_dir, f"confusion_matrix_{clf.replace(' ', '_')}.csv")
    if os.path.exists(cm_file):
        cm_df = pd.read_csv(cm_file, index_col=0)

        plt.figure(figsize=(6.5, 5.5))
        sns.heatmap(cm_df, annot=True, fmt='g', cbar=True)
        plt.title(f"Confusion Matrix — {clf}")
        plt.ylabel('True label')
        plt.xlabel('Predicted label')
        plt.tight_layout()

        out_path = os.path.join(out_dir, f"confusion_matrix_{clf.replace(' ', '_')}_heatmap.png")
        plt.savefig(out_path, dpi=200, bbox_inches="tight")
        plt.close()

print("Confusion matrix heatmaps (if any) have been saved.")
print(f"Summary CSV saved to: {summary_csv_path}")
```
