## Supplementary material for "Determinants of Blood Group Antigen Expression and Prediction of Phenotypes by Machine Learning": ML Supplement: boxplot_visualisations_with_stat_sign_a.py

```
import os
import sys
import numpy as np
import pandas as pd
import matplotlib.pyplot as plt
import seaborn as sns

# out_dir comes from CLI: python boxplot_visualisations_with_stat_sign.py <output_dir>
out_dir = sys.argv[1] if len(sys.argv) > 1 else "."
file_path = os.path.join(out_dir, "fold_results.csv")

os.makedirs(out_dir, exist_ok=True)

# Matplotlib font size
plt.rcParams.update({'font.size': 20})

# =========================
# Load
# =========================
df = pd.read_csv(file_path)

# =========================
# NEW: Load pairwise significance results (if available)
# =========================
sig_dir = os.path.join(out_dir, "significance_tests")
pairwise_sig_path = os.path.join(sig_dir, "pairwise_significance.csv")

if os.path.exists(pairwise_sig_path):
    sig_df = pd.read_csv(pairwise_sig_path)
else:
    sig_df = None
    print("No pairwise_significance.csv found -> boxplots will not show significance annotations.")

# =========================
# Helpers for significance annotation
# =========================
def p_to_stars(p):
    """Convert p-value to significance stars."""
    if p <= 0.001:
        return "***"
    elif p <= 0.01:
        return "**"
    elif p <= 0.05:
        return "*"
    else:
        return "ns"

def get_metric_name_for_sig(metric_vis_name):
    """
    Map the metric name used in the fold_results.csv / plots
    to the 'Metric' name stored in pairwise_significance.csv.
    Adjust this mapping to match **your** training script if needed.
    """
    mapping = {
        # If your training script uses these:
        "Test F1-Macro": "Test F1-Macro",
        "Test Balanced Accuracy": "Test Balanced Accuracy",

        # If your CSV uses the alternative wording:
        "F1-Macro on test data": "F1-Macro on test data",
        "Balanced Accuracy on test data": "Balanced Accuracy on test data",
    }
    return mapping.get(metric_vis_name, metric_vis_name)

def annotate_significance_for_metric(ax, df, metric, sig_df, max_pairs=10):
    """
    Add significance bars on top of a boxplot, using pairwise_significance.csv.

    ax     : matplotlib Axes with the boxplot
    df     : per-fold results DataFrame (contains 'Classifier' and metric columns)
    metric : metric column name as used in df / plotting
    sig_df : full significance DataFrame (loaded from CSV)
    """
    if sig_df is None:
        return

    # Only annotate for macro F1 and balanced accuracy
    metrics_of_interest = {
        "F1-Macro on test data",
        "Test F1-Macro",
        "Balanced Accuracy on test data",
        "Test Balanced Accuracy",
    }
    if metric not in metrics_of_interest:
        return

    metric_sig_name = get_metric_name_for_sig(metric)
    metric_sig_df = sig_df[sig_df["Metric"] == metric_sig_name].copy()
    if metric_sig_df.empty:
        print(f"No significance info found for metric '{metric_sig_name}' in pairwise_significance.csv")
        return

    # Keep only significant pairs (Holm-adjusted p < 0.05)
    if "p_value_adj_holm" in metric_sig_df.columns:
        metric_sig_df = metric_sig_df[metric_sig_df["p_value_adj_holm"] < 0.05]
        metric_sig_df = metric_sig_df.sort_values("p_value_adj_holm")
    elif "Significant(p<0.05)" in metric_sig_df.columns:
        metric_sig_df = metric_sig_df[metric_sig_df["Significant(p<0.05)"] == True]
        metric_sig_df = metric_sig_df.sort_values("p_value_raw")
    else:
        print(f"pairwise_significance.csv has no adjusted p-values / significance flag; skipping annotations for {metric}.")
        return

    if metric_sig_df.empty:
        print(f"No significant pairs (Holm-adjusted) for metric '{metric_sig_name}'")
        return

    # Limit how many pairs we annotate to avoid total chaos
    metric_sig_df = metric_sig_df.head(max_pairs)

    # Map classifier name -> x position in the boxplot
    xticklabels = [tick.get_text() for tick in ax.get_xticklabels()]
    x_positions = {clf: i for i, clf in enumerate(xticklabels)}

    # Base height for the first bar
    data_max = df[metric].max()
    y_start = data_max + 0.02
    h = 0.03  # vertical spacing between bars

    # Draw the lines
    for idx, row in metric_sig_df.iterrows():
        clf_a = row["Model_A"]
        clf_b = row["Model_B"]

        if clf_a not in x_positions or clf_b not in x_positions:
            # Might happen if some classifiers were filtered out in the plot
            continue

        x1 = x_positions[clf_a]
        x2 = x_positions[clf_b]
        if x1 == x2:
            continue
        if x1 > x2:
            x1, x2 = x2, x1

        # height for this pair
        y = y_start + (idx * h)

        # Get p-value (adjusted if present)
        p = row["p_value_adj_holm"] if "p_value_adj_holm" in row else row["p_value_raw"]
        stars = p_to_stars(p)

        # Draw bar
        ax.plot([x1, x1, x2, x2],
                [y, y + 0.01, y + 0.01, y],
                color="black", linewidth=1.0)

        # Add text
        ax.text((x1 + x2) / 2.0, y + 0.012, stars,
                ha="center", va="bottom", color="black", fontsize=12)

    # Increase ylimit so annotations are visible
    ymin, ymax = ax.get_ylim()
    ax.set_ylim(ymin, max(ymax, y_start + (len(metric_sig_df) + 1) * h))


# =========================
# Choose metrics dynamically (handles old & new CSVs)
# =========================
all_candidate_metrics = [
    "Train Accuracy", "Test Accuracy", "Test Precision", "Test Recall", "Test F1-Score",
    "F1-Macro on test data", "Balanced Accuracy on test data",
    ### NEW: also support the names from your training script directly
    "Test F1-Macro", "Test Balanced Accuracy"
]
metrics = [m for m in all_candidate_metrics if m in df.columns]

if not metrics:
    raise ValueError("No known metric columns found in the CSV. "
                     "Expected one of: " + ", ".join(all_candidate_metrics))

# =========================
# Boxplots per metric (with significance for macro F1 and balanced accuracy)
# =========================
for metric in metrics:
    plt.figure(figsize=(10, 6))
    ax = sns.boxplot(x="Classifier", y=metric, data=df, hue="Classifier", palette="Set3")
    plt.title(f"{metric}")
    plt.ylabel(metric)
    plt.xlabel("Classifier")
    plt.xticks(rotation=45, ha="right")
    plt.legend([], [], frameon=False)  # hide legend (duplicate of x labels)
    plt.subplots_adjust(bottom=0.35)   # more space for rotated labels

    # Default y-limits for probabilities
    ax.set_ylim(0, 1)

    # === NEW: add significance bars for macro F1 and balanced accuracy (if available) ===
    annotate_significance_for_metric(ax, df, metric, sig_df, max_pairs=10)

    plot_file_path = os.path.join(out_dir, f"{metric.replace(' ', '_')}_boxplot.png")
    plt.savefig(plot_file_path, dpi=200, bbox_inches="tight")
    plt.close()

print("Box plots have been saved successfully.")

# =========================
# Summary table: mean & std per metric/classifier
# =========================
if metrics:
    agg_spec = {m: ['mean', 'std'] for m in metrics}
    summary_data = df.groupby('Classifier').agg(agg_spec).reset_index()

    # Flatten MultiIndex columns: ("Test Accuracy","mean") -> "Test Accuracy Mean"
    new_cols = []
    for c in summary_data.columns:
        if isinstance(c, tuple):
            if c[1] == '':
                new_cols.append(c[0])  # e.g. "Classifier"
            else:
                new_cols.append(f"{c[0]} {'Mean' if c[1] == 'mean' else 'Std Dev'}")
        else:
            new_cols.append(c)
    summary_data.columns = new_cols

else:
    raise ValueError("No known metric columns found in the CSV. ")

# Save summary table
summary_csv_path = os.path.join(out_dir, "metrics_summary_by_classifier.csv")
summary_data.to_csv(summary_csv_path, index=False)

# =========================
# Bar charts (mean ± std) per metric
# =========================
classifiers = summary_data['Classifier']
plt.rcParams.update({'font.size': 22})

for metric in metrics:
    mean_col = f"{metric} Mean"
    std_col  = f"{metric} Std Dev"

    if mean_col not in summary_data.columns:
        continue

    means = summary_data[mean_col]
    std_devs = summary_data[std_col] if std_col in summary_data.columns else None

    x_pos = np.arange(len(classifiers))

    plt.figure(figsize=(12, 8))
    plt.bar(x_pos, means, yerr=std_devs, align='center', alpha=0.7, ecolor='black', capsize=11)
    plt.xticks(x_pos, classifiers, rotation=45, ha='right')
    plt.xlabel('Classifier')
    plt.ylabel(metric)
    plt.title(f'{metric} Across Classifiers')
    plt.ylim(0, 1)
    plt.tight_layout()

    plot_file_path = os.path.join(out_dir, f"{metric.replace(' ', '_')}_bar_chart.png")
    plt.savefig(plot_file_path, dpi=200, bbox_inches="tight")
    plt.close()

print("Bar charts have been saved successfully.")

# =========================
# OPTIONAL: Confusion matrix heatmaps (if CSVs exist)
# =========================
unique_classifiers = df['Classifier'].unique()
for clf in unique_classifiers:
    cm_file = os.path.join(out_dir, f"confusion_matrix_{clf.replace(' ', '_')}.csv")
    if os.path.exists(cm_file):
        cm_df = pd.read_csv(cm_file, index_col=0)

        plt.figure(figsize=(6.5, 5.5))
        sns.heatmap(cm_df, annot=True, fmt='g', cbar=True)
        plt.title(f"Confusion Matrix — {clf}")
        plt.ylabel('True label')
        plt.xlabel('Predicted label')
        plt.tight_layout()

        out_path = os.path.join(out_dir, f"confusion_matrix_{clf.replace(' ', '_')}_heatmap.png")
        plt.savefig(out_path, dpi=200, bbox_inches="tight")
        plt.close()

print("Confusion matrix heatmaps (if any) have been saved.")
print(f"Summary CSV saved to: {summary_csv_path}")
```
