## Supplementary material for "Determinants of Blood Group Antigen Expression and Prediction of Phenotypes by Machine Learning": ML Supplement: feature_importance_visualization_a.py

```
import os
import sys
import pandas as pd
import numpy as np
import matplotlib.pyplot as plt

# out_dir comes from CLI: python feature_importance_visualization.py <output_dir>
out_dir = sys.argv[1] if len(sys.argv) > 1 else "."
csv_path = os.path.join(out_dir, "feature_importances_variance_summary.csv")
df = pd.read_csv(csv_path)

# Ensure we have a standard deviation column; compute it from variance if needed
if "std" not in df.columns and "var" in df.columns:
    df["std"] = np.sqrt(df["var"])
elif "std" not in df.columns:
    raise ValueError("CSV must contain either 'std' or 'var' to draw error bars.")

classifiers = df["Classifier"].unique()
features = df["Feature"].unique()

x = np.arange(len(features))
width = 0.8 / len(classifiers)

plt.figure(figsize=(12, 6))

for i, clf in enumerate(classifiers):
    subset = df[df["Classifier"] == clf]
    subset = subset.set_index("Feature").reindex(features).reset_index()

    plt.bar(
        x + i * width,
        subset["mean"].values,
        yerr=subset["std"].values,
        width=width,
        label=clf,
        capsize=5
    )

plt.title("Mean Feature Importance per Classifier")
plt.xlabel("Feature")
plt.ylabel("Mean Importance (± SD)")
plt.xticks(x + width * (len(classifiers) - 1) / 2, features, rotation=45, ha="right")
plt.legend(title="Classifier")
plt.tight_layout()

plt.savefig(os.path.join(out_dir, "feature_importances_grouped_errorbars.png"), dpi=300)
```
