## Supplementary material for "Determinants of Blood Group Antigen Expression and Prediction of Phenotypes by Machine Learning": ML Supplement: train_classifier_12_05_26.py

```
# train_delta_features_ctrlv_nullv_grouped_with_significance.py
# Group-aware nested CV (keeps all samples from the same blood group together)
# + Statistical significance testing across classifiers
# + Nested grouped repeated CV: outer 5x3, inner 2
# + Generalized repeated 5x3 paired t/F tests on grouped outer-split differences
#
# NOTE:
# - Canonical Dietterich test is 5x2cv paired t-test.
# - Canonical Alpaydin test is combined 5x2cv F-test.
# - This script implements a grouped repeated-k-fold analogue for 5x3
#   because your requested design is 5 seeds x 3 folds.

import os
import json
import re
import sys
import subprocess
import warnings

import numpy as np
import pandas as pd
import matplotlib

matplotlib.use("Agg")
import matplotlib.pyplot as plt

from pathlib import Path
from imblearn.under_sampling import RandomUnderSampler
from imblearn.pipeline import Pipeline as ImbPipeline

from sklearn import __version__ as sklearn_version
from sklearn.impute import SimpleImputer
from sklearn.preprocessing import StandardScaler
from sklearn.dummy import DummyClassifier
from sklearn.linear_model import LogisticRegression
from sklearn.ensemble import RandomForestClassifier, GradientBoostingClassifier
from sklearn.metrics import (
    make_scorer, accuracy_score, precision_score, recall_score,
    f1_score, balanced_accuracy_score, confusion_matrix, classification_report
)
from sklearn.model_selection import GridSearchCV
from sklearn.svm import LinearSVC
from sklearn.calibration import CalibratedClassifierCV
from sklearn.inspection import permutation_importance
from sklearn.exceptions import UndefinedMetricWarning

with warnings.catch_warnings():
    warnings.filterwarnings(
        "ignore",
        message="y_pred contains classes not in y_true",
        category=UserWarning,
    )
    warnings.filterwarnings(
        "ignore",
        category=UndefinedMetricWarning,
    )

try:
    from scipy import stats

    SCIPY_OK = True
except Exception as e:
    SCIPY_OK = False
    print("Warning: SciPy not available; significance tests will be skipped. Error:", e)


def collapse_phenotype(phenotype: str) -> str:
    if phenotype in ["AgV", "AgWT"]:
        return "Antigenic"
    elif phenotype == "CtrlV":
        return "CtrlV"
    else:
        return "Other"


# ======================================================
# GLOBAL CONFIG
# ======================================================
USE_GPT = False
file_path = "../merged_variant_data_78_a.csv"

TASKS_TO_RUN = [
    "CTRLV_VS_AG",
    "CTRLV_VS_NULLV",
]

OUTER_N_SPLITS = 3
OUTER_SEEDS = [11, 22, 33, 44, 55]  # 5 repeats / seeds
INNER_N_SPLITS = 2
INNER_SEED = 42

BASE_OUTPUT_ROOT = f"example_outputs/train_features_grouped_repeated_5x{OUTER_N_SPLITS}x{INNER_N_SPLITS}_011"
os.makedirs(BASE_OUTPUT_ROOT, exist_ok=True)

# ======================================================
# FEATURES & TARGET
# ======================================================
FEATURE_SETS = {
    "CTRLV_VS_NULLV": [
        "Delta_Hydrophobicity",
        "Delta_Sidechain Volume",
        "SurfaceAccessibility",
        "MedusaFlexWeighted",
        "Conservation",
        "MissenseScore",
    ],
    "CTRLV_VS_AG": [
        "Hydrophobicity",
        "Sidechain Volume",
        "SurfaceAccessibility",
        "Conservation",
        "MedusaFlexWeighted",
    ],
}


def run_task(TASK):
    print("\n" + "=" * 80)
    print(f"RUNNING TASK: {TASK}")
    print("=" * 80)

    output_dir = os.path.join(
        BASE_OUTPUT_ROOT,
        f"train_features_{TASK.lower()}_results_grouped_repeated_5x{OUTER_N_SPLITS}x{INNER_N_SPLITS}_v01"
    )
    os.makedirs(output_dir, exist_ok=True)

    log_file = os.path.join(output_dir, "cross_validation_results.txt")
    sig_dir = os.path.join(output_dir, "significance_tests")
    os.makedirs(sig_dir, exist_ok=True)

    features = FEATURE_SETS[TASK]
    FEATURE_NAMES = np.array(features)

    target = "Phenotype"
    GROUP_COL = "BloodGroup"

    # ======================================================
    # LOAD DATA
    # ======================================================
    protein_data = pd.read_csv(file_path)

    client = None # OpenAI(api_key="sk-0000")

    if TASK == "CTRLV_VS_NULLV":
        df = protein_data[protein_data["Phenotype"].isin(["CtrlV", "NullV"])].copy()
        target = "Phenotype"
        df[target] = pd.Categorical(df[target], categories=["CtrlV", "NullV"], ordered=True)

    elif TASK == "CTRLV_VS_AG":
        df = protein_data[protein_data["Phenotype"].isin(["CtrlV", "AgV", "AgWT"])].copy()
        df["Phenotype_bin"] = df["Phenotype"].replace({"AgV": "Ag", "AgWT": "Ag"})
        target = "Phenotype_bin"
        df[target] = pd.Categorical(df[target], categories=["CtrlV", "Ag"], ordered=True)

    else:
        raise ValueError(f"Unknown TASK='{TASK}'. Use 'CTRLV_VS_NULLV' or 'CTRLV_VS_AG'.")

    missing = [c for c in features + [target, GROUP_COL] if c not in df.columns]
    if missing:
        raise ValueError(f"Missing required columns: {missing}")

    for c in features:
        df[c] = pd.to_numeric(df[c], errors="coerce")

    df = df.dropna(subset=[target]).reset_index(drop=True)

    class_names = df[target].cat.categories
    y = df[target].cat.codes.values
    ALL_LABELS = np.arange(len(class_names))
    X = df[features].values
    groups = df[GROUP_COL].astype("category")

    print("Class distribution:")
    print(df[target].value_counts(dropna=False))
    print("\n# unique groups:", groups.nunique())

    # ======================================================
    # GPT HELPERS
    # ======================================================
    def build_few_shot_examples_balanced_from_df(train_fold_df, n_per_class=3, seed=42):
        rng = np.random.RandomState(seed)

        parts = []
        for cls in train_fold_df[target].cat.categories:
            pool = train_fold_df[
                (train_fold_df[target] == cls) &
                train_fold_df[features].notna().all(axis=1)
                ]
            if len(pool) == 0:
                continue

            k = min(n_per_class, len(pool))
            few = pool.sample(n=k, random_state=rng.randint(0, 10000), replace=False)

            for _, row in few.iterrows():
                desc = "\n".join([f"{f}: {row[f]}" for f in features])
                parts.append(f"{desc}\nPhenotype: {cls}")

        if len(parts) == 0:
            raise ValueError("No few-shot examples available in this fold.")

        return "\n\n".join(parts)

    def gpt_classify_batch(rows_df, few_shot_examples, class_names, id_col="__rid__"):
        cases = []
        for _, r in rows_df.iterrows():
            case = {"id": str(r[id_col])}
            for f in features:
                v = r[f]
                case[f] = None if pd.isna(v) else float(v)
            cases.append(case)

        prompt = make_gpt_prompt(
            cases=cases,
            few_shot_examples=few_shot_examples,
            class_names=class_names,
            features=features,
        )

        resp = client.chat.completions.create(
            model="gpt-5.4-2026-03-05",
            messages=[{"role": "user", "content": prompt}],
        )
        text = resp.choices[0].message.content

        obj = None
        try:
            m = re.search(r"\{.*\}", text, re.S)
            if m:
                obj = json.loads(m.group(0))
        except Exception:
            obj = None

        parsed = parse_gpt_predictions_object(
            obj=obj,
            class_names=class_names,
            features=features,
        )
        return parsed

    # ======================================================
    # CV SPLITTERS (GROUP-AWARE)
    # ======================================================
    try:
        from sklearn.model_selection import StratifiedGroupKFold

        HAS_SGF = True
    except Exception:
        StratifiedGroupKFold = None
        HAS_SGF = False

    def make_outer_splitter(seed):
        if HAS_SGF:
            return StratifiedGroupKFold(
                n_splits=OUTER_N_SPLITS,
                shuffle=True,
                random_state=seed,
            )
        else:
            from sklearn.model_selection import GroupKFold
            return GroupKFold(n_splits=OUTER_N_SPLITS)

    def make_inner_splitter():
        if HAS_SGF:
            return StratifiedGroupKFold(
                n_splits=INNER_N_SPLITS,
                shuffle=True,
                random_state=INNER_SEED,
            )
        else:
            from sklearn.model_selection import GroupKFold
            return GroupKFold(n_splits=INNER_N_SPLITS)

    inner_cv = make_inner_splitter()

    if HAS_SGF:
        print(f"Using StratifiedGroupKFold: outer={len(OUTER_SEEDS)}x{OUTER_N_SPLITS}, inner={INNER_N_SPLITS}.")
    else:
        print(
            f"StratifiedGroupKFold not available in sklearn {sklearn_version}; "
            f"using GroupKFold for grouped outer={len(OUTER_SEEDS)}x{OUTER_N_SPLITS} and inner={INNER_N_SPLITS}."
        )

    # ======================================================
    # PIPELINE FACTORY
    # ======================================================
    def make_pipeline(clf, do_undersample=False, strategy="not minority", random_state=42):
        steps = [("imputer", SimpleImputer(strategy="mean"))]
        if do_undersample:
            steps.append(("undersample", RandomUnderSampler(sampling_strategy=strategy, random_state=random_state)))
        steps += [("scaler", StandardScaler()), ("clf", clf)]
        return ImbPipeline(steps)

    # ======================================================
    # CLASSIFIERS & PARAM GRIDS
    # ======================================================
    classifiers = {
        "Dummy (Most Frequent)": make_pipeline(
            DummyClassifier(strategy="most_frequent", random_state=42),
            do_undersample=False
        ),
        "Logistic Regression": make_pipeline(
            LogisticRegression(max_iter=1000, random_state=42),
            do_undersample=False
        ),
        "Random Forest": make_pipeline(
            RandomForestClassifier(n_estimators=100, random_state=42),
            do_undersample=False
        ),
        "Gradient Boosting": make_pipeline(
            GradientBoostingClassifier(random_state=42),
            do_undersample=False
        ),
        "Support Vector Machine": make_pipeline(
            LinearSVC(random_state=42),
            do_undersample=False
        ),
    }

    param_grids = {
        "Dummy (Most Frequent)": {},
        "Logistic Regression": {
            "clf__C": [0.001, 0.01, 0.1, 1],  # "clf__C": [0.01, 0.1, 1, 10, 100],
            "clf__solver": ["liblinear", "lbfgs"],
            "clf__penalty": ["l2"],
            "clf__class_weight": [None, "balanced"],
        },
        "Random Forest": {
            "clf__n_estimators": [200, 500],
            "clf__max_depth": [3, 5, 8],  # 10
            "clf__min_samples_split": [5, 10, 20],
            "clf__min_samples_leaf": [2, 4, 8],
            "clf__max_features": ["sqrt"],
            "clf__class_weight": ["balanced", "balanced_subsample"],
        },
        # "Random Forest": {
        #    "clf__n_estimators": [200, 500],
        #    "clf__max_depth": [None, 5, 10, 20],
        #    "clf__min_samples_split": [2, 5, 10],
        #    "clf__min_samples_leaf": [1, 2, 4],
        #    "clf__max_features": ["sqrt", "log2", None],
        #    "clf__class_weight": [None, "balanced", "balanced_subsample"],
        # },
        # "Gradient Boosting": {
        #    "clf__n_estimators": [100, 200, 500],
        #    "clf__learning_rate": [0.01, 0.05, 0.1, 0.2],
        #    "clf__max_depth": [1, 2, 3],
        #    "clf__subsample": [0.6, 0.8, 1.0],
        # },
        "Gradient Boosting": {
            "clf__n_estimators": [50, 100, 200],
            "clf__learning_rate": [0.01, 0.05],
            # "clf__n_estimators": [100, 200],
            # "clf__learning_rate": [0.01, 0.05, 0.1],
            "clf__max_depth": [1, 2],
            "clf__subsample": [0.6, 0.8],
        },
        "Support Vector Machine": {
            #"clf__C": [0.001, 0.01, 0.1, 1],
            "clf__C": [0.01, 0.1, 1, 10, 100],
            # "clf__estimator__C": [0.01, 0.1, 1, 10, 100],
            "clf__class_weight": [None, "balanced"],
            # "clf__estimator__class_weight": [None, "balanced"],
        },
    }

    scoring = {
        "Accuracy": make_scorer(accuracy_score),
        "Precision": make_scorer(precision_score, average="weighted", zero_division=1, labels=ALL_LABELS),
        "Recall": make_scorer(recall_score, average="weighted", zero_division=1, labels=ALL_LABELS),
        "F1": make_scorer(f1_score, average="weighted", zero_division=1, labels=ALL_LABELS),
        "F1_macro": make_scorer(f1_score, average="macro", zero_division=1, labels=ALL_LABELS),
        "Balanced_Accuracy": make_scorer(balanced_accuracy_score),
    }

    # ======================================================
    # MAIN TRAIN / EVAL
    # ======================================================
    results_overall = {}
    fold_results = []
    best_params_per_fold = []
    feature_importances_across_folds = []

    with open(log_file, "w") as log:
        for name, base_pipe in classifiers.items():
            print(f"\nNested repeated grouped CV (tuning) for {name}...")
            log.write(f"\nNested repeated grouped CV (tuning) for {name}...\n")

            # --------------------------------------------------
            # Dummy baseline (manual repeated grouped outer loop)
            # --------------------------------------------------
            if name == "Dummy (Most Frequent)":
                split_counter = 0
                y_pred_dummy_all = pd.Series(index=range(len(y)), dtype=float)
                acc_tr, f1m_tr, bal_tr, acc_te, prec_te, rec_te, f1_te, f1m_te, bal_te = ([] for _ in range(9))

                for repeat_idx, outer_seed in enumerate(OUTER_SEEDS, start=1):
                    outer_cv = make_outer_splitter(outer_seed)

                    for fold_idx, (tr_idx, te_idx) in enumerate(outer_cv.split(X, y, groups), start=1):
                        split_counter += 1

                        X_tr, X_te = X[tr_idx], X[te_idx]
                        y_tr, y_te = y[tr_idx], y[te_idx]

                        est = base_pipe.fit(X_tr, y_tr)
                        y_tr_pred = est.predict(X_tr)
                        y_te_pred = est.predict(X_te)

                        for idx_pos, pred in zip(te_idx, y_te_pred):
                            if pd.isna(y_pred_dummy_all.iloc[idx_pos]):
                                y_pred_dummy_all.iloc[idx_pos] = pred

                        tr_acc = accuracy_score(y_tr, y_tr_pred)
                        tr_f1m = f1_score(y_tr, y_tr_pred, average="macro", zero_division=1, labels=ALL_LABELS)
                        tr_bal = balanced_accuracy_score(y_tr, y_tr_pred)

                        te_acc = accuracy_score(y_te, y_te_pred)
                        te_prec = precision_score(y_te, y_te_pred, average="weighted", zero_division=1,
                                                  labels=ALL_LABELS)
                        te_rec = recall_score(y_te, y_te_pred, average="weighted", zero_division=1, labels=ALL_LABELS)
                        te_f1 = f1_score(y_te, y_te_pred, average="weighted", zero_division=1, labels=ALL_LABELS)
                        te_f1m = f1_score(y_te, y_te_pred, average="macro", zero_division=1, labels=ALL_LABELS)
                        te_bal = balanced_accuracy_score(y_te, y_te_pred)

                        fold_results.append({
                            "Classifier": name,
                            "Repeat": repeat_idx,
                            "Seed": outer_seed,
                            "Fold": fold_idx,
                            "SplitID": split_counter,
                            "Train Accuracy": tr_acc,
                            "Train F1-Macro": tr_f1m,
                            "Train Balanced Accuracy": tr_bal,
                            "Test Accuracy": te_acc,
                            "Test Precision": te_prec,
                            "Test Recall": te_rec,
                            "Test F1-Score": te_f1,
                            "Test F1-Macro": te_f1m,
                            "Test Balanced Accuracy": te_bal,
                        })

                        log.write(
                            f"[{name}] Repeat {repeat_idx} Seed {outer_seed} Fold {fold_idx} - "
                            f"Train Acc: {tr_acc:.4f}, Test Acc: {te_acc:.4f}, "
                            f"Prec: {te_prec:.4f}, Rec: {te_rec:.4f}, F1: {te_f1:.4f}, "
                            f"F1-macro: {te_f1m:.4f}, BalAcc: {te_bal:.4f}\n"
                        )

                        acc_tr.append(tr_acc)
                        f1m_tr.append(tr_f1m)
                        bal_tr.append(tr_bal)
                        acc_te.append(te_acc)
                        prec_te.append(te_prec)
                        rec_te.append(te_rec)
                        f1_te.append(te_f1)
                        f1m_te.append(te_f1m)
                        bal_te.append(te_bal)

                results_overall[name] = {
                    "Train Accuracy": np.mean(acc_tr),
                    "Train F1-Macro": np.mean(f1m_tr),
                    "Train Balanced Accuracy": np.mean(bal_tr),
                    "Test Accuracy": np.mean(acc_te),
                    "Test Precision": np.mean(prec_te),
                    "Test Recall": np.mean(rec_te),
                    "Test F1-Score": np.mean(f1_te),
                    "Test F1-Macro": np.mean(f1m_te),
                    "Test Balanced Accuracy": np.mean(bal_te),
                }

                filled = y_pred_dummy_all.fillna(0).astype(int).values
                cls_rep = classification_report(
                    y, filled,
                    target_names=list(class_names),
                    output_dict=True,
                    zero_division=1
                )
                pd.DataFrame(cls_rep).transpose().to_csv(
                    os.path.join(output_dir, f"per_class_report_{name.replace(' ', '_')}.csv"),
                    index=True
                )
                cm = confusion_matrix(y, filled)
                pd.DataFrame(cm, index=class_names, columns=class_names).to_csv(
                    os.path.join(output_dir, f"confusion_matrix_{name.replace(' ', '_')}.csv")
                )

                continue

            # --------------------------------------------------
            # Tuned models: manual repeated grouped outer loop
            # --------------------------------------------------
            grid = param_grids[name]
            refit_score = "F1_macro"

            outer_predictions_all = {}

            acc_tr, f1m_tr, bal_tr, acc_te, prec_te, rec_te, f1_te, f1m_te, bal_te = ([] for _ in range(9))
            split_counter = 0

            for repeat_idx, outer_seed in enumerate(OUTER_SEEDS, start=1):
                outer_cv = make_outer_splitter(outer_seed)

                for fold_idx, (tr_idx, te_idx) in enumerate(outer_cv.split(X, y, groups), start=1):
                    split_counter += 1

                    X_tr, X_te = X[tr_idx], X[te_idx]
                    y_tr, y_te = y[tr_idx], y[te_idx]
                    groups_tr = groups.iloc[tr_idx] if hasattr(groups, "iloc") else groups[tr_idx]

                    gs = GridSearchCV(
                        estimator=base_pipe,
                        param_grid=grid,
                        scoring=scoring,
                        refit=refit_score,
                        cv=inner_cv,
                        n_jobs=-1,
                    )
                    gs.fit(X_tr, y_tr, groups=groups_tr)

                    if hasattr(gs.best_estimator_, "named_steps") and "undersample" in gs.best_estimator_.named_steps:
                        sampler = gs.best_estimator_.named_steps["undersample"]
                        _, y_tr_bal = sampler.fit_resample(X_tr, y_tr)
                        unique, counts = np.unique(y_tr_bal, return_counts=True)
                        print(f"[Fold {fold_idx}] Balanced train counts:", dict(zip(unique, counts)))

                    best_est = gs.best_estimator_
                    best_params = gs.best_params_

                    best_params_per_fold.append({
                        "Classifier": name,
                        "Repeat": repeat_idx,
                        "Seed": outer_seed,
                        "Fold": fold_idx,
                        "SplitID": split_counter,
                        **best_params
                    })

                    # === Capture per-fold feature importances ===
                    try:
                        clf_step = best_est.named_steps["clf"]

                        if hasattr(clf_step, "feature_importances_"):
                            imp = clf_step.feature_importances_
                            denom = np.sum(imp)
                            imp_norm = imp / (denom if denom != 0 else 1.0)

                            fi_fold = pd.DataFrame({
                                "Classifier": name,
                                "Repeat": repeat_idx,
                                "Seed": outer_seed,
                                "Fold": fold_idx,
                                "SplitID": split_counter,
                                "Feature": features,
                                "Importance": imp_norm,
                            })
                            feature_importances_across_folds.append(fi_fold)

                        elif hasattr(clf_step, "coef_"):
                            coef_abs = np.mean(np.abs(clf_step.coef_), axis=0)
                            denom = np.sum(coef_abs)
                            coef_abs_norm = coef_abs / (denom if denom != 0 else 1.0)

                            fi_fold = pd.DataFrame({
                                "Classifier": name,
                                "Repeat": repeat_idx,
                                "Seed": outer_seed,
                                "Fold": fold_idx,
                                "SplitID": split_counter,
                                "Feature": features,
                                "Importance": coef_abs_norm,
                            })
                            feature_importances_across_folds.append(fi_fold)

                    except Exception as e:
                        print(f"[Fold {fold_idx}] Skipped feature importance capture for {name}: {e}")

                    y_tr_pred = best_est.predict(X_tr)
                    y_te_pred = best_est.predict(X_te)

                    for idx_pos, pred in zip(te_idx, y_te_pred):
                        if idx_pos not in outer_predictions_all:
                            outer_predictions_all[idx_pos] = pred

                    tr_acc = accuracy_score(y_tr, y_tr_pred)
                    tr_f1m = f1_score(y_tr, y_tr_pred, average="macro", zero_division=1, labels=ALL_LABELS)
                    tr_bal = balanced_accuracy_score(y_tr, y_tr_pred)

                    te_acc = accuracy_score(y_te, y_te_pred)
                    te_prec = precision_score(y_te, y_te_pred, average="weighted", zero_division=1, labels=ALL_LABELS)
                    te_rec = recall_score(y_te, y_te_pred, average="weighted", zero_division=1, labels=ALL_LABELS)
                    te_f1 = f1_score(y_te, y_te_pred, average="weighted", zero_division=1, labels=ALL_LABELS)
                    te_f1m = f1_score(y_te, y_te_pred, average="macro", zero_division=1, labels=ALL_LABELS)
                    te_bal = balanced_accuracy_score(y_te, y_te_pred)

                    fold_results.append({
                        "Classifier": name,
                        "Repeat": repeat_idx,
                        "Seed": outer_seed,
                        "Fold": fold_idx,
                        "SplitID": split_counter,
                        "Train Accuracy": tr_acc,
                        "Train F1-Macro": tr_f1m,
                        "Train Balanced Accuracy": tr_bal,
                        "Test Accuracy": te_acc,
                        "Test Precision": te_prec,
                        "Test Recall": te_rec,
                        "Test F1-Score": te_f1,
                        "Test F1-Macro": te_f1m,
                        "Test Balanced Accuracy": te_bal,
                    })

                    log.write(
                        f"[{name}] Repeat {repeat_idx} Seed {outer_seed} Fold {fold_idx} "
                        f"(SplitID={split_counter}) best params: {best_params}\n"
                    )
                    log.write(
                        f"[{name}] Repeat {repeat_idx} Seed {outer_seed} Fold {fold_idx} - "
                        f"Train Acc: {tr_acc:.4f}, Test Acc: {te_acc:.4f}, "
                        f"Prec: {te_prec:.4f}, Rec: {te_rec:.4f}, F1: {te_f1:.4f}, "
                        f"F1-macro: {te_f1m:.4f}, BalAcc: {te_bal:.4f}\n"
                    )

                    acc_tr.append(tr_acc)
                    f1m_tr.append(tr_f1m)
                    bal_tr.append(tr_bal)
                    acc_te.append(te_acc)

                    prec_te.append(te_prec)
                    rec_te.append(te_rec)
                    f1_te.append(te_f1)
                    f1m_te.append(te_f1m)
                    bal_te.append(te_bal)

            results_overall[name] = {
                "Train Accuracy": np.mean(acc_tr),
                "Train F1-Macro": np.mean(f1m_tr),
                "Train Balanced Accuracy": np.mean(bal_tr),
                "Test Accuracy": np.mean(acc_te),
                "Test Precision": np.mean(prec_te),
                "Test Recall": np.mean(rec_te),
                "Test F1-Score": np.mean(f1_te),
                "Test F1-Macro": np.mean(f1m_te),
                "Test Balanced Accuracy": np.mean(bal_te),
            }

            log.write(f"Overall (repeated grouped outer-CV) results for {name}:\n")
            for k, v in results_overall[name].items():
                log.write(f"  {k}: {v:.4f}\n")

            y_pred_all = np.full(shape=len(y), fill_value=-1, dtype=int)
            for idx_pos, pred in outer_predictions_all.items():
                y_pred_all[idx_pos] = pred
            y_pred_all[y_pred_all == -1] = 0

            cls_rep = classification_report(
                y,
                y_pred_all,
                labels=list(ALL_LABELS),
                target_names=list(class_names),
                output_dict=True,
                zero_division=1
            )
            pd.DataFrame(cls_rep).transpose().to_csv(
                os.path.join(output_dir, f"per_class_report_{name.replace(' ', '_')}.csv"),
                index=True
            )
            cm = confusion_matrix(y, y_pred_all)
            pd.DataFrame(cm, index=class_names, columns=class_names).to_csv(
                os.path.join(output_dir, f"confusion_matrix_{name.replace(' ', '_')}.csv")
            )

    # ======================================================
    # GPT-5 BASELINE ON THE SAME OUTER SPLITS
    # ======================================================
    if USE_GPT:
        from openai import OpenAI
        from gpt_posthoc_utils import (
            make_gpt_prompt,
            parse_gpt_predictions_object,
            save_gpt_posthoc_outputs,
        )
        gpt_name = "GPT-5 (few-shot)"

        acc_te, prec_te, rec_te, f1_te, f1m_te, bal_te = ([] for _ in range(6))
        gpt_fold_prediction_rows = []

        split_counter = 0

        for repeat_idx, outer_seed in enumerate(OUTER_SEEDS, start=1):
            outer_cv = make_outer_splitter(outer_seed)

            for fold_idx, (tr_idx, te_idx) in enumerate(outer_cv.split(X, y, groups), start=1):
                split_counter += 1
                print(f"\n[{gpt_name}] Repeat {repeat_idx} Seed {outer_seed} Fold {fold_idx}...")

                train_fold_df = df.iloc[tr_idx].copy()
                test_fold_df = df.iloc[te_idx].copy()

                few_shot = build_few_shot_examples_balanced_from_df(
                    train_fold_df,
                    n_per_class=3,
                    seed=42 + repeat_idx + fold_idx
                )

                BATCH_SIZE = 12
                test_fold_df = test_fold_df.copy()
                test_fold_df["__rid__"] = test_fold_df.index.astype(str)

                pred_map = {}
                n_rows = len(test_fold_df)
                n_batches = int(np.ceil(n_rows / BATCH_SIZE))

                for b in range(n_batches):
                    start = b * BATCH_SIZE
                    end = min((b + 1) * BATCH_SIZE, n_rows)
                    batch_df = test_fold_df.iloc[start:end]

                    print(
                        f"[{gpt_name}] Repeat {repeat_idx} Fold {fold_idx}: "
                        f"batch {b + 1}/{n_batches} ({end - start} rows)"
                    )
                    batch_preds = gpt_classify_batch(batch_df, few_shot, class_names, id_col="__rid__")
                    pred_map.update(batch_preds)

                y_true = y[te_idx]
                y_pred_codes = []

                for _, row in test_fold_df.iterrows():
                    rid = str(row["__rid__"])
                    pred_obj = pred_map.get(rid, {})
                    lab = pred_obj.get("Phenotype", "Unknown")

                    if lab in list(class_names):
                        y_pred_codes.append(list(class_names).index(lab))
                    else:
                        y_pred_codes.append(0)

                    gpt_fold_prediction_rows.append({
                        "Repeat": repeat_idx,
                        "Seed": outer_seed,
                        "Fold": fold_idx,
                        "SplitID": split_counter,
                        "__rid__": rid,
                        target: row[target],
                        "GPT5_JSON": json.dumps(pred_obj.get("raw_prediction", pred_obj)),
                        "PredictedLabel": lab,
                        "Confidence": pred_obj.get("confidence", np.nan),
                    })

                y_pred_codes = np.array(y_pred_codes, dtype=int)

                te_acc = accuracy_score(y_true, y_pred_codes)
                te_prec = precision_score(y_true, y_pred_codes, average="weighted", zero_division=1, labels=ALL_LABELS)
                te_rec = recall_score(y_true, y_pred_codes, average="weighted", zero_division=1, labels=ALL_LABELS)
                te_f1 = f1_score(y_true, y_pred_codes, average="weighted", zero_division=1, labels=ALL_LABELS)
                te_f1m = f1_score(y_true, y_pred_codes, average="macro", zero_division=1, labels=ALL_LABELS)
                te_bal = balanced_accuracy_score(y_true, y_pred_codes)

                fold_results.append({
                    "Classifier": gpt_name,
                    "Repeat": repeat_idx,
                    "Seed": outer_seed,
                    "Fold": fold_idx,
                    "SplitID": split_counter,
                    "Train Accuracy": np.nan,
                    "Test Accuracy": te_acc,
                    "Test Precision": te_prec,
                    "Test Recall": te_rec,
                    "Test F1-Score": te_f1,
                    "Test F1-Macro": te_f1m,
                    "Test Balanced Accuracy": te_bal,
                })

                acc_te.append(te_acc)
                prec_te.append(te_prec)
                rec_te.append(te_rec)
                f1_te.append(te_f1)
                f1m_te.append(te_f1m)
                bal_te.append(te_bal)

        results_overall[gpt_name] = {
            "Train Accuracy": np.nan,
            "Test Accuracy": np.mean(acc_te),
            "Test Precision": np.mean(prec_te),
            "Test Recall": np.mean(rec_te),
            "Test F1-Score": np.mean(f1_te),
            "Test F1-Macro": np.mean(f1m_te),
            "Test Balanced Accuracy": np.mean(bal_te),
        }
    else:
        gpt_fold_prediction_rows = []

    # ======================================================
    # SAVE SUMMARIES
    # ======================================================
    best_params_df = pd.DataFrame(best_params_per_fold)
    best_params_df.to_csv(os.path.join(output_dir, "best_params_per_fold.csv"), index=False)

    fold_results_df = pd.DataFrame(fold_results)
    results_df = pd.DataFrame(results_overall).T

    fold_results_df.to_csv(os.path.join(output_dir, "fold_results.csv"), index=False)
    results_df.to_csv(os.path.join(output_dir, "cross_validation_results.csv"), index=True)

    if USE_GPT:
        gpt_fold_predictions_df = pd.DataFrame(gpt_fold_prediction_rows)
        gpt_fold_predictions_df.to_csv(
            os.path.join(output_dir, "gpt_fold_predictions_raw.csv"),
            index=False
        )
    else:
        gpt_fold_predictions_df = pd.DataFrame()

    # ======================================================
    # GPT POST-HOC FEATURE IMPORTANCE
    # ======================================================
    gpt_posthoc_dir = os.path.join(output_dir, "gpt_posthoc")
    os.makedirs(gpt_posthoc_dir, exist_ok=True)

    if not gpt_fold_predictions_df.empty:
        gpt_posthoc_outputs = save_gpt_posthoc_outputs(
            gpt_fold_predictions_df=gpt_fold_predictions_df,
            features=features,
            output_dir=gpt_posthoc_dir,
            target_col=target,
            gpt_name=gpt_name,
            fold_results_df=fold_results_df,
        )
        if gpt_posthoc_outputs["tf_long_df"]["Importance"].sum() == 0:
            raise RuntimeError("GPT posthoc feature importance is all zero; check top_features parsing.")
        print("Saved GPT posthoc outputs to:", gpt_posthoc_dir)
    else:
        print("No GPT raw predictions available for posthoc analysis.")

    # ======================================================
    # AGGREGATE PER-FOLD FEATURE IMPORTANCES
    # ======================================================
    if feature_importances_across_folds:
        fi_df = pd.concat(feature_importances_across_folds, ignore_index=True)
        fi_path = os.path.join(output_dir, "feature_importances_per_fold.csv")
        fi_df.to_csv(fi_path, index=False)

        fi_var = (
            fi_df.groupby(["Classifier", "Feature"])["Importance"]
            .agg(mean="mean", var="var", std="std", count="count")
            .reset_index()
        )
        fi_var_path = os.path.join(output_dir, "feature_importances_variance_summary.csv")
        fi_var.to_csv(fi_var_path, index=False)
        print("Saved per-fold importances to:", fi_path)
        print("Saved variance summary to:", fi_var_path)

    print("\nPer-Fold Results:")
    print(fold_results_df.head())
    print("\nOverall Cross-Validation Results:")
    print(results_df)

    # ======================================================
    # SIGNIFICANCE TESTS
    # ======================================================
    def holm_bonferroni(pvals):
        m = len(pvals)
        order = np.argsort(pvals)
        adj = np.empty(m, dtype=float)
        prev = 0.0
        for k, i in enumerate(order):
            adj_i = (m - k) * pvals[i]
            adj[i] = max(prev, adj_i)
            prev = adj[i]
        return np.minimum(adj, 1.0)

    def build_repeat_fold_diff_matrix(
            fold_results_df,
            metric,
            model_a,
            model_b,
            expected_repeats=5,
            expected_folds=3
    ):
        sub = fold_results_df[fold_results_df["Classifier"].isin([model_a, model_b])].copy()

        piv = sub.pivot_table(
            index=["Repeat", "Fold"],
            columns="Classifier",
            values=metric,
            aggfunc="mean"
        ).dropna()

        needed = {model_a, model_b}
        if not needed.issubset(set(piv.columns)):
            return None

        piv = piv[[model_a, model_b]].copy()
        piv["diff"] = piv[model_a] - piv[model_b]

        counts = piv.reset_index().groupby("Repeat")["Fold"].nunique()
        valid_repeats = counts[counts == expected_folds].index.tolist()

        piv = piv.reset_index()
        piv = piv[piv["Repeat"].isin(valid_repeats)]

        if len(valid_repeats) != expected_repeats:
            return None

        piv = piv.sort_values(["Repeat", "Fold"])
        D = piv.pivot(index="Repeat", columns="Fold", values="diff").values

        if D.shape != (expected_repeats, expected_folds):
            return None

        return D

    def dietterich_repeated_kfold_t_approx(D):
        """
        Generalized repeated-k-fold paired t analogue.
        D shape = (R, K), here expected (5, 3).
        This is NOT the canonical Dietterich 5x2cv t-test.
        """
        R, K = D.shape
        d_bar = D.mean()

        s2 = np.var(D, axis=1, ddof=1)
        s2_mean = np.mean(s2)

        se = np.sqrt(s2_mean / (R * K) + 1e-12)
        t_stat = d_bar / se

        dfree = R * (K - 1)
        p_val = 2.0 * stats.t.sf(np.abs(t_stat), df=dfree)

        return {
            "t_stat": float(t_stat),
            "df": int(dfree),
            "p_value": float(p_val),
            "mean_diff": float(d_bar),
            "within_repeat_var_mean": float(s2_mean),
        }

    def alpaydin_repeated_kfold_f_approx(D):
        """
        Generalized repeated-k-fold F analogue.
        D shape = (R, K), here expected (5, 3).
        This is NOT the canonical Alpaydin 5x2cv combined F-test.
        """
        R, K = D.shape

        num = np.sum(D ** 2)
        repeat_means = D.mean(axis=1, keepdims=True)
        denom_ss = np.sum((D - repeat_means) ** 2)

        df1 = R * K
        df2 = R * (K - 1)

        if denom_ss <= 1e-12:
            F_stat = np.inf if num > 0 else 0.0
            p_val = 0.0 if np.isfinite(F_stat) and F_stat > 0 else 1.0
        else:
            F_stat = (num / df1) / (denom_ss / df2)
            p_val = stats.f.sf(F_stat, df1, df2)

        return {
            "F_stat": float(F_stat),
            "df1": int(df1),
            "df2": int(df2),
            "p_value": float(p_val),
            "sum_sq_diff": float(num),
            "within_repeat_ss": float(denom_ss),
        }

    def alpaydin_repeated_kfold_f_approx(D):
        """
        Generalized repeated-k-fold F analogue.
        D shape = (R, K), here expected (5, 3).
        This is NOT the canonical Alpaydin 5x2cv combined F-test.
        """
        R, K = D.shape

        num = np.sum(D ** 2)
        repeat_means = D.mean(axis=1, keepdims=True)
        denom_ss = np.sum((D - repeat_means) ** 2)

        df1 = R * K
        df2 = R * (K - 1)

        if denom_ss <= 1e-12:
            F_stat = np.inf if num > 0 else 0.0
            p_val = 0.0 if np.isfinite(F_stat) and F_stat > 0 else 1.0
        else:
            F_stat = (num / df1) / (denom_ss / df2)
            p_val = stats.f.sf(F_stat, df1, df2)

        return {
            "F_stat": float(F_stat),
            "df1": int(df1),
            "df2": int(df2),
            "p_value": float(p_val),
            "sum_sq_diff": float(num),
            "within_repeat_ss": float(denom_ss),
        }

    if SCIPY_OK and not fold_results_df.empty:
        metrics_to_test = [
            "Test Accuracy",
            "Test F1-Score",
            "Test Balanced Accuracy",
            "Test F1-Macro",
        ]

        from itertools import combinations

        repeated_kfold_rows = []

        for metric in metrics_to_test:
            clfs = sorted(fold_results_df["Classifier"].unique().tolist())

            for model_a, model_b in combinations(clfs, 2):
                D = build_repeat_fold_diff_matrix(
                    fold_results_df=fold_results_df,
                    metric=metric,
                    model_a=model_a,
                    model_b=model_b,
                    expected_repeats=len(OUTER_SEEDS),
                    expected_folds=OUTER_N_SPLITS,
                )

                if D is None:
                    print(
                        f"[Repeated-k-fold tests] Skipping {metric}: "
                        f"{model_a} vs {model_b} (incomplete repeat/fold grid)"
                    )
                    continue

                try:
                    t_res = dietterich_repeated_kfold_t_approx(D)
                    f_res = alpaydin_repeated_kfold_f_approx(D)
                except Exception as e:
                    print(f"[Significance] Failed for {metric}, {model_a} vs {model_b}: {e}")
                    continue

                repeated_kfold_rows.append({
                    "Metric": metric,
                    "Model_A": model_a,
                    "Model_B": model_b,
                    "Test": "Dietterich_like_repeated_5x3_t",
                    "Statistic": t_res["t_stat"],
                    "df1": t_res["df"],
                    "df2": np.nan,
                    "p_value_raw": t_res["p_value"],
                    "mean_diff(A-B)": t_res["mean_diff"],
                    "within_repeat_var_mean": t_res["within_repeat_var_mean"],
                    "n_repeats": D.shape[0],
                    "n_folds_per_repeat": D.shape[1],
                })

                repeated_kfold_rows.append({
                    "Metric": metric,
                    "Model_A": model_a,
                    "Model_B": model_b,
                    "Test": "Alpaydin_like_repeated_5x3_F",
                    "Statistic": f_res["F_stat"],
                    "df1": f_res["df1"],
                    "df2": f_res["df2"],
                    "p_value_raw": f_res["p_value"],
                    "mean_diff(A-B)": float(D.mean()),
                    "within_repeat_var_mean": float(np.mean(np.var(D, axis=1, ddof=1))),
                    "n_repeats": D.shape[0],
                    "n_folds_per_repeat": D.shape[1],
                })

        if repeated_kfold_rows:
            repeated_kfold_df = pd.DataFrame(repeated_kfold_rows)

            out_blocks = []
            for (metric, test_name), block in repeated_kfold_df.groupby(["Metric", "Test"], dropna=False):
                block = block.copy()
                block["p_value_adj_holm"] = holm_bonferroni(block["p_value_raw"].values)
                block["Significant(p<0.05)"] = block["p_value_adj_holm"] < 0.05
                out_blocks.append(block)

            repeated_kfold_df = pd.concat(out_blocks, ignore_index=True)
            repeated_kfold_df.sort_values(["Metric", "Test", "p_value_adj_holm"], inplace=True)
            repeated_kfold_df.to_csv(
                os.path.join(sig_dir, "pairwise_repeated_5x3_grouped_significance.csv"),
                index=False
            )
            print("Saved repeated grouped 5x3 significance results.")
    else:
        if not SCIPY_OK:
            print("SciPy missing -> skipping significance tests. Install with `pip install scipy`.")
        else:
            print("No fold_results available for significance testing.")

    # ======================================================
    # FIT BEST RF / GB / LR / SVM ON FULL DATA
    # ======================================================
    def preprocess_from_pipeline(fitted_pipe, X_raw):
        X_imp = fitted_pipe.named_steps["imputer"].transform(X_raw)
        X_scl = fitted_pipe.named_steps["scaler"].transform(X_imp)
        return X_scl

    # --- Random Forest ---
    rf_pipe = classifiers["Random Forest"]
    rf_grid = param_grids["Random Forest"]
    rf_gs = GridSearchCV(
        estimator=rf_pipe,
        param_grid=rf_grid,
        scoring=scoring,
        refit="F1_macro",
        cv=inner_cv,
        n_jobs=-1
    )
    rf_gs.fit(X, y, groups=groups)
    best_rf = rf_gs.best_estimator_
    print(f"\nBest RF params (full-data search): {rf_gs.best_params_}")

    rf_model = best_rf.named_steps["clf"]
    X_rf_proc = preprocess_from_pipeline(best_rf, X)
    rf_importances = rf_model.feature_importances_
    rf_imp_df = pd.DataFrame({"Feature": features, "Importance": rf_importances}).sort_values("Importance",
                                                                                              ascending=False)
    rf_imp_df.to_csv(os.path.join(output_dir, "feature_importance_rf.csv"), index=False)
    """
    X_rf_proc = np.asarray(X_rf_proc, dtype=float)
    expl_rf = shap.TreeExplainer(rf_model)
    sv_rf = expl_rf.shap_values(X_rf_proc)
    sv_list = sv_rf if isinstance(sv_rf, list) else [sv_rf]
    rf_bee = np.mean(np.abs(np.stack(sv_list, axis=0)), axis=0)

    plt.figure(figsize=(8, 5))
    shap.summary_plot(rf_bee, feature_names=FEATURE_NAMES, plot_type="bar", show=False)
    plt.title("SHAP Feature Importance (RF, mean |SHAP| across classes)")
    plt.tight_layout()
    plt.savefig(os.path.join(output_dir, "shap_rf_bar.png"), dpi=300)
    plt.close()

    plt.figure(figsize=(8, 5))
    shap.summary_plot(rf_bee, X_rf_proc, feature_names=FEATURE_NAMES, show=False)
    plt.title("SHAP Beeswarm (RF, mean |SHAP| across classes)")
    plt.tight_layout()
    plt.savefig(os.path.join(output_dir, "shap_rf_beeswarm.png"), dpi=300)
    plt.close()
"""
    # --- Gradient Boosting ---
    gb_pipe = classifiers["Gradient Boosting"]
    gb_grid = param_grids["Gradient Boosting"]
    gb_gs = GridSearchCV(
        estimator=gb_pipe,
        param_grid=gb_grid,
        scoring=scoring,
        refit="F1_macro",
        cv=inner_cv,
        n_jobs=-1
    )
    gb_gs.fit(X, y, groups=groups)
    best_gb = gb_gs.best_estimator_
    print(f"\nBest GB params (full-data search): {gb_gs.best_params_}")

    gb_model = best_gb.named_steps["clf"]
    X_gb_proc = preprocess_from_pipeline(best_gb, X)
    X_gb_proc = np.asarray(X_gb_proc, dtype=float)

    gb_importances = gb_model.feature_importances_
    gb_imp_df = pd.DataFrame({"Feature": features, "Importance": gb_importances}).sort_values("Importance",
                                                                                              ascending=False)
    gb_imp_df.to_csv(os.path.join(output_dir, "feature_importance_gb.csv"), index=False)

    expl_gb = shap.TreeExplainer(gb_model)
    sv_gb = expl_gb.shap_values(X_gb_proc)
    sv_list_gb = sv_gb if isinstance(sv_gb, list) else [sv_gb]
    gb_bee = np.mean(np.abs(np.stack(sv_list_gb, axis=0)), axis=0)

    plt.figure(figsize=(8, 5))
    shap.summary_plot(gb_bee, feature_names=FEATURE_NAMES, plot_type="bar", show=False)
    plt.title("SHAP Feature Importance (GB, mean |SHAP| across classes)")
    plt.tight_layout()
    plt.savefig(os.path.join(output_dir, "shap_gb_bar.png"), dpi=300)
    plt.close()

    plt.figure(figsize=(8, 5))
    shap.summary_plot(gb_bee, X_gb_proc, feature_names=FEATURE_NAMES, show=False)
    plt.title("SHAP Beeswarm (GB, mean |SHAP| across classes)")
    plt.tight_layout()
    plt.savefig(os.path.join(output_dir, "shap_gb_beeswarm.png"), dpi=300)
    plt.close()

    # --- Logistic Regression ---
    lr_pipe = classifiers["Logistic Regression"]
    lr_grid = param_grids["Logistic Regression"]
    lr_gs = GridSearchCV(
        estimator=lr_pipe,
        param_grid=lr_grid,
        scoring=scoring,
        refit="F1_macro",
        cv=inner_cv,
        n_jobs=-1
    )
    lr_gs.fit(X, y, groups=groups)
    best_lr = lr_gs.best_estimator_
    print(f"\nBest LR params (full-data search): {lr_gs.best_params_}")

    lr_model = best_lr.named_steps["clf"]
    X_lr_proc = preprocess_from_pipeline(best_lr, X)
    X_lr_proc = np.asarray(X_lr_proc, dtype=float)

    coef_abs = np.mean(np.abs(lr_model.coef_), axis=0)
    denom = coef_abs.sum()
    coef_abs_norm = coef_abs / (denom if denom != 0 else 1.0)

    lr_imp_df = (
        pd.DataFrame({"Feature": features, "Importance": coef_abs_norm})
        .sort_values("Importance", ascending=False)
    )
    lr_imp_df.to_csv(os.path.join(output_dir, "feature_importance_lr_coef_abs.csv"), index=False)

    expl_lr = shap.LinearExplainer(lr_model, X_lr_proc)
    sv_lr = expl_lr.shap_values(X_lr_proc)
    sv_list_lr = sv_lr if isinstance(sv_lr, list) else [sv_lr]
    lr_bee = np.mean(np.abs(np.stack(sv_list_lr, axis=0)), axis=0)

    plt.figure(figsize=(8, 5))
    shap.summary_plot(lr_bee, feature_names=FEATURE_NAMES, plot_type="bar", show=False)
    plt.title("SHAP Feature Importance (LR, mean |SHAP| across classes)")
    plt.tight_layout()
    plt.savefig(os.path.join(output_dir, "shap_lr_bar.png"), dpi=300)
    plt.close()

    plt.figure(figsize=(8, 5))
    shap.summary_plot(lr_bee, X_lr_proc, feature_names=FEATURE_NAMES, show=False)
    plt.title("SHAP Beeswarm (LR, mean |SHAP| across classes)")
    plt.tight_layout()
    plt.savefig(os.path.join(output_dir, "shap_lr_beeswarm.png"), dpi=300)
    plt.close()

    perm = permutation_importance(
        best_lr, X, y,
        scoring="f1_macro",
        n_repeats=20,
        random_state=42,
        n_jobs=-1
    )
    perm_df = pd.DataFrame({"Feature": features, "Importance": perm.importances_mean}).sort_values("Importance",
                                                                                                   ascending=False)
    perm_df.to_csv(os.path.join(output_dir, "feature_importance_lr_permutation.csv"), index=False)

    # --- SVM (LinearSVC + calibration) ---
    svm_pipe = classifiers["Support Vector Machine"]
    svm_grid = param_grids["Support Vector Machine"]
    svm_gs = GridSearchCV(
        estimator=svm_pipe,
        param_grid=svm_grid,
        scoring=scoring,
        refit="F1_macro",
        cv=inner_cv,
        n_jobs=-1
    )
    svm_gs.fit(X, y, groups=groups)
    best_svm = svm_gs.best_estimator_
    print(f"\nBest SVM params (full-data search): {svm_gs.best_params_}")

    svm_lin = best_svm.named_steps["clf"]
    X_svm_proc = preprocess_from_pipeline(best_svm, X)

    coef_abs = np.mean(np.abs(svm_lin.coef_), axis=0)
    denom = coef_abs.sum()
    coef_abs_norm = coef_abs / (denom if denom != 0 else 1.0)

    svm_imp_df = (
        pd.DataFrame({"Feature": features, "Importance": coef_abs_norm})
        .sort_values("Importance", ascending=False)
    )
    svm_imp_df.to_csv(os.path.join(output_dir, "feature_importance_svm_coef_abs.csv"), index=False)

    # Comparison table
    rf_imp_df = pd.read_csv(os.path.join(output_dir, "feature_importance_rf.csv"))
    gb_imp_df = pd.read_csv(os.path.join(output_dir, "feature_importance_gb.csv"))

    cmp = (
        lr_imp_df.rename(columns={"Importance": "LR_coef_abs"})
        .merge(rf_imp_df.rename(columns={"Importance": "RF_imp"}), on="Feature", how="outer")
        .merge(gb_imp_df.rename(columns={"Importance": "GB_imp"}), on="Feature", how="outer")
        .fillna(0.0)
    )

    for col in ["LR_coef_abs", "RF_imp", "GB_imp"]:
        s = cmp[col].to_numpy()
        denom = s.sum() if s.sum() != 0 else 1.0
        cmp[col + "_norm"] = s / denom

    cmp = cmp.sort_values("LR_coef_abs_norm", ascending=False)
    cmp.to_csv(os.path.join(output_dir, "feature_importance_comparison_lr_rf_gb.csv"), index=False)

    print("\nSaved outputs to:", output_dir)

    # ======================================================
    # POST-HOC VISUALISATIONS
    # ======================================================
    scripts_dir = Path(__file__).resolve().parent
    subprocess.run([sys.executable, scripts_dir / "feature_importance_visualization_a.py", output_dir], check=True)
    boxplot_script = scripts_dir / "boxplot_visualisations_with_stat_sign_a.py"

    subprocess.run(
        [sys.executable, str(boxplot_script), str(output_dir)],
        check=True
    )


if __name__ == "__main__":
    for task_name in TASKS_TO_RUN:
        run_task(task_name)
```
